# Mixing and matching methylotrophic enzymes to design a novel methanol utilization pathway in *E. coli*

**DOI:** 10.1101/2020.04.15.042333

**Authors:** A. De Simone, C.M. Vicente, C. Peiro, L. Gales, F. Bellvert, B. Enjalbert, S. Heux

**Affiliations:** TBI, Université de Toulouse, CNRS, INRAE, INSA, Toulouse, France; MetaboHUB-MetaToul, National Infrastructure of Metabolomics and Fluxomics, Toulouse, 31077, France

**Keywords:** One-carbon metabolism, Methanol, *Escherichia coli*, Synthetic methylotrophy

## Abstract

One-carbon (C1) compounds, such as methanol, have recently gained attention as alternative low-cost and non-food feedstocks for microbial bioprocesses. Considerable research efforts are thus currently focused on the generation of synthetic methylotrophs by transferring methanol assimilation pathways into established bacterial production hosts. In this study, we used an iterative combination of dry and wet approaches to design, implement and optimize this metabolic trait in the most common chassis, *E. coli*. Through *in silico* modeling, we designed a new route that “mixed and matched” two methylotrophic enzymes: a bacterial methanol dehydrogenase (Mdh) and a dihydroxyacetone synthase (Das) from yeast. To identify the best combination of enzymes to introduce into *E. coli*, we built a library of 266 pathway variants containing different combinations of Mdh and Das homologues and screened it using high-throughput ^13^C-labeling experiments. The highest level of incorporation, 22% of labeled methanol carbon into the multi-carbon compound PEP, was obtained using a variant composed of a Mdh from *A. gerneri* and a codon-optimized version of *P. angusta* Das. Finally, the activity of this new synthetic pathway was further improved by engineering strategic metabolic targets identified using omics and modelling approaches. The final synthetic strain had 1.5 to 5.9 times higher methanol assimilation in intracellular metabolites and proteinogenic amino acids than the starting strain did. Broadening the repertoire of methanol assimilation pathways is one step further toward synthetic methylotrophy in *E. coli*.

## 1. INTRODUCTION

In the quest to replace fossil fuel-based processes with more sustainable bio-based ones, low-cost and easy to use fermentation substrates are of great interest. Commonly used feedstocks such as hydrolyzed starch and molasses have the disadvantage of competing with food supply, and lignocellulosic biomass requires costly pre-treatment. A promising alternative feedstock is methanol, an abundant and pure raw material that can be utilized directly in bacterial fermentation processes. Methanol’s higher degree of reduction means furthermore that it is more energy rich than carbohydrates and this extra energy can be expected to enhance product yields during fermentation. Methanol is currently one of the top five commodity chemicals with a global production capacity of about 110 million metric tons per year and a price similar to that of glucose (http://www.methanol.org/). Although methanol is mainly produced from fossil resources, a notable advantage is that it can be produced by polygeneration, as a product of any renewable resource that can be converted into an intermediated synthesis gas (syngas). This includes biomass, agricultural and timber waste, municipal solid waste, landfill gas, industrial waste and a number of other feedstocks (http://enerkem.com/fr/; http://www.methanol.org/). Bio-methanol can also be produced by chemically recycling methane and CO2, two notorious greenhouse gases (http://www.biomcn.eu/; http://www.carbonrecycling.is/). All these factors make methanol an attractive feedstock for biorefineries and the concept of a methanol economy has received considerable attention (Olah, 2013; Schrader et al., 2009).

Methylotrophy is the capacity of certain prokaryote and eukaryote microorganisms to use reduced one-carbon (C1) compounds such as methanol or methane as their sole source of carbon and energy. This metabolism includes: (i) the oxidation of methanol to formaldehyde; (ii) the oxidation of formaldehyde to CO_2_, and (iii) the assimilation of one carbon compounds, either formaldehyde or CO2 or a combination thereof (Heux S. et al., 2018). The industrial-scale use of natural methylotrophs has already been attempted. In the 1970s, a process was developed to produce single-cell protein (SCP) from methanol (Matelbs and Tannenbaum, 1968; Windass et al., 1980), but the technology fell out of favor in the following decades because of the low prices of alternative sources such as soybean protein. Nowadays, the use of natural methylotrophs in bioprocesses is only seen in the production by methylotrophic yeasts of recombinant proteins such as enzymes, antibodies, cytokines, plasma proteins, and hormones (Ahmad et al., 2014). The production of small molecules and metabolites (e.g. PHAs (polyhydroxyalkanoates) and amino acids) is still at the proof-of-concept stage (Schrader et al., 2009). The main limitations to the use of natural methylotrophs in biotechnologies are our currently weak understanding of their cellular metabolism and physiology, and the general lack of genetic tools to modify them (Chung et al., 2010; Schrader et al., 2009). In contrast, *Escherichia coli* is a robust biotechnological chassis with a wide range of products and an extensive genetic toolbox (Becker and Wittmann, 2015). Engineering a methanol assimilation pathway in this microorganism has thus become a popular research topic.

Methylotrophy is quite challenging to engineer because all biomass production and energy requirements must be satisfied by a reduced C1 precursor. In addition, cells must be able to tolerate formaldehyde, a central but toxic compound in methanol metabolism, whose accumulation due to an imbalance between oxidation and assimilation in the pathway can be fatal for cells. Because formaldehyde oxidation is efficient, the main bottleneck is C1 assimilation, which is achieved through a cyclic process involving a C1-acceptor to enable the formation of C-C bonds. Several attempts have been made to engineer synthetic methylotrophy in *E. coli* using naturally occurring cyclic pathways (Wang et al., 2020). Most of these involve the expression of three heterologous enzymes: a NAD^+^-dependent methanol dehydrogenase (Mdh) for the oxidation of methanol to formaldehyde together with hexulose phosphate synthase (Hps) and phosphohexuloisomerase (Phi) from the ribulose monophosphate (RuMP) cycle for formaldehyde fixation. The *in vivo* operation of this pathway in *E. coli* has been confirmed by isotope-labeling experiments, which showed that methanol carbons were incorporated into cellular material (Muller et al., 2015). Similar results have also been reported in other model organisms such as *Corynebacterium glutamicum, Pseudomonas putida* and *Saccharomyces cerevisiae* (as reviewed recently by (Heux S. et al., 2018)). Improvements in methanol assimilation have been achieved using different strategies such as (i) optimizing the cultivation medium (Gonzalez et al., 2018), (ii) lowering the thermodynamic and kinetic constraints associated with NAD-dependent methanol oxidation (Roth et al., 2019; Wu et al., 2016), (iii) improving formaldehyde assimilation (Price et al., 2016; Woolston et al., 2018), (iv) increasing carbon fluxes through the autocatalytic cycle (Bennett et al., 2018), and (v) coupling the activity of the RuMP cycle to the growth of the host microorganism and then using adaptive laboratory evolution (Chen et al., 2018; He et al., 2018; Meyer et al., 2018). However, none of these synthetic strains are able to grow on methanol alone. The reasons for this and the obstacles to overcome include regenerating the C1-acceptor, protecting the cells against formaldehyde toxicity, channeling the substrate so that it can be integrated directly into the central metabolism, and lowering energetic constraints.

The approach outlined here to tackle the exciting challenge of synthetic methylotrophy is to develop a hybrid of naturally occurring cyclic methanol assimilation pathways. Using a “mix and match” approach, we created a new synthetic pathway combining Mdh, a methylotrophic enzyme of bacterial origin, with dihydroxyacetone synthase (Das), a methylotrophic enzyme from yeast. The engineered strain was then optimized in an iterative process using omics (transcriptomics, metabolomics and fluxomics) and modelling approaches to identify bottlenecks. Overall, this approach allows non-natural pathways to be explored and tested while offering new perspectives on synthetic methylotrophy in *E. coli*.

## 2. RESULTS

### 2.1. Selecting the best design for a methanol assimilation pathway

Natural methylotrophs have developed multiple pathways that allow them to grow on methanol as the sole source of carbon and energy (Chistoserdova, 2011). This metabolic diversity means that in theory, there are more than 1000 unique methanol assimilation pathways from methanol to biomass. To identify the best pathway for *E. coli* to consume methanol, we used FindPath, a tool that freely recombines a repertoire of existing reactions to create metabolic pathways (Vieira et al., 2014). FindPath uses a substrate-associated reaction database and flux balance analysis (FBA) based on a genome scale model (GSM) of the host to (i) find all the possible pathways, and (ii) rank them according to their length and the predicted growth rate on the substrate of interest. The tool identified two equally efficient synthetic routes: the already well-studied RuMP-based pathway involving the bacterial enzymes Mdh, Hps and Phi, and a hybrid metabolic pathway, involving methylotrophic bacterial Mdh and methylotrophic yeast derived Das (Figure 1). The latter is a transketolase that catalyzes the fixation of formaldehyde on xylulose 5-phosphate (Xu5P) to form glyceraldehyde 3-phosphate (GAP) and dihydroxyacetone (DHA) in the xylulose monophosphate (XuMP) cycle in methylotrophic yeasts. The GSM-predicted growth rate of *E. coli* on methanol with this pathway is 0.34 h^−1^. The predicted fluxes show that this optimal growth rate is achieved when 16% of the methanol is incorporated into the biomass with the rest being used to recycle the C1 acceptor, Xu5P. No flux through the methanol oxidation pathway (i.e. through FrmA & B) was predicted. In addition, the GSM predicted that Xu5P would be recycled by fructose-6-phosphate aldolase and transaldolase (FSA/TAL pathway variant) rather than by fructose 1,6-bisphosphate aldolase with transaldolase (FBA/TAL variant), or by sedoheptulose biphosphatase (FBA/SBP variant) (Supplementary Figure S1). No ATP is required for Xu5P regeneration in the FSA/TAL variant, while the other two metabolic variants require two ATP molecules (Supplementary Figure S1). In comparison, in the RuMP based pathway, the C1 acceptor, ribulose-5-phosphate, is recycled using one or two ATP molecules (Woolston et al., 2018). This and the fact that our synthetic pathway involves the expression of two enzymes instead of the three as in the RuMP based-pathway, indicates that, in theory, the novel route is more energetically favorable without any need to dissimilate methanol into formate (He et al., 2020). In addition, the transketolase activity of Das may contribute to the regeneration of the C1 acceptor, making our pathway partially independent of the pentose phosphate pathway (PPP). This has been demonstrated recently in methylotrophic yeasts (Russmayer et al., 2015).

**Figure 1.**
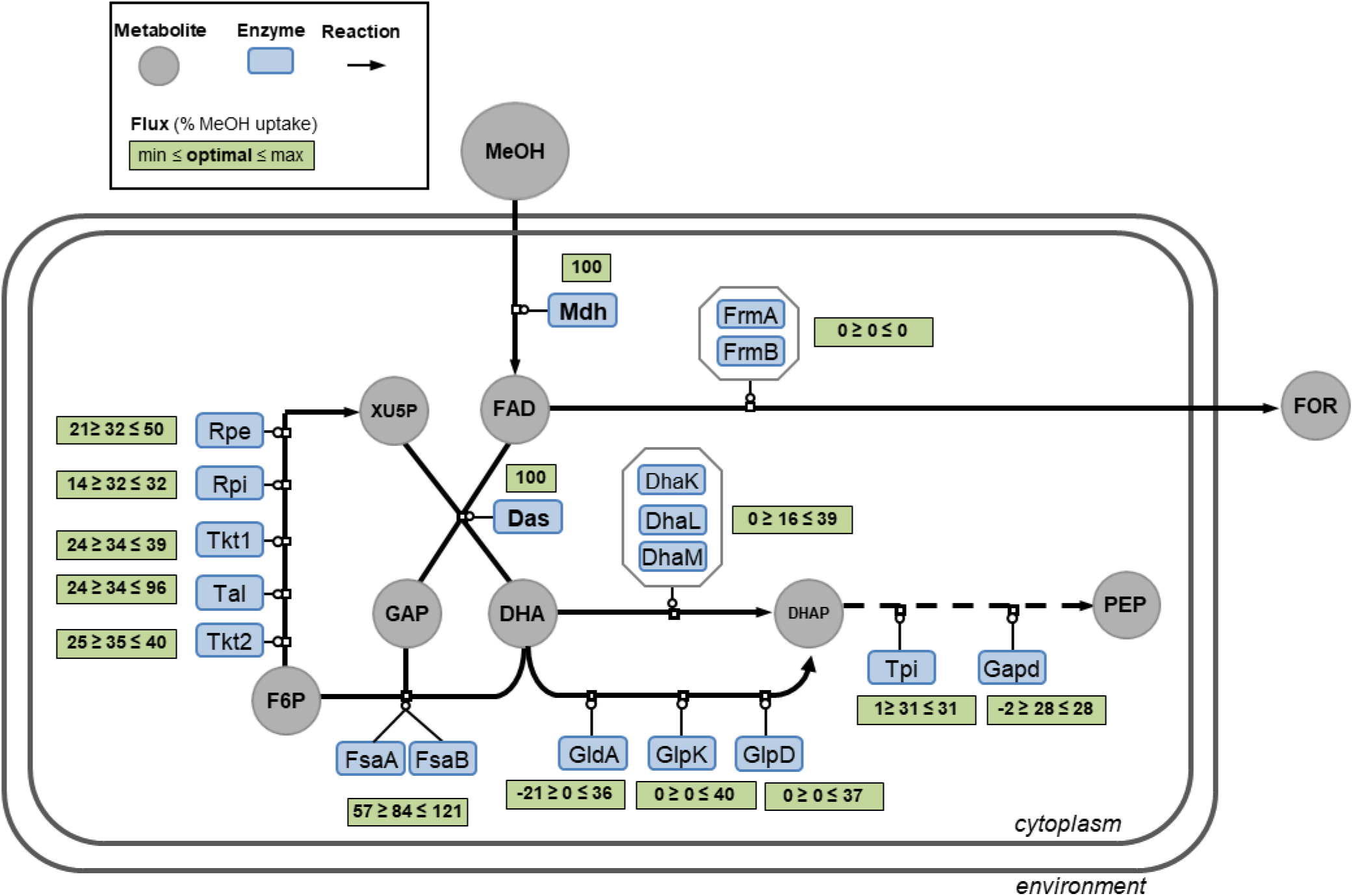
Overview of the synthetic methanol metabolism and its operation in *E. coli*. The new hybrid methanol assimilation pathway comprises a methanol dehydrogenase (Mdh) and a dihydroxyacetone synthase (Das). Green rectangles give the optimal and the ranges of simulated fluxes obtained using flux balance analysis and flux variability analysis, respectively, when growth rate is constrained to 90% of the optimal value. Flux values are given in % relative to a MeOH uptake rate of 15 mmol/g_DW_/h as defined in Peyraud et al., BMC Syst Biol. 2011. Dihydroxyacetone kinase (DhaK, DhaL and DhaM); Glycerol dehydrogenase (GldA); Glycerol-3-phosphate dehydrogenase (GlpD); Glycerol kinase (GlpK); Fructose-6-phosphate aldolase (FsaA and FsaB); Triose phosphate isomerase (Tpi); Glyceraldehyde-3-phosphate dehydrogenase (Gapd); Ribulose 5-phosphate 3-epimerase (Rpe); Transketolase (Tkt1 & Tkt2); Transaldolase (Tal), Ribose-5-phosphate isomerase (Rpi); Formaldehyde dehydrogenase (FrmA); S-Formylglutathione hydrolase (FrmB); Methanol (MeOH); Formaldehyde (FAD); Xylulose-5-P (XU5P); Glyceraldehyde-3-phosphate (GAP); Dihydroxyacetone (DHA); Phosphoenolpyruvate (PEP); Dihydroxyacetone phosphate (DHAP); Fructose-6-phosphate (F6P), Formate (FOR).

### 2.2. Screening for the best matching methylotrophic enzymes

To identify the combination of enzymes that would optimize the *in vivo* activity of the pathway in *E. coli*, we built a combinatorial library of *mdh* and *das* homologues derived from native and non-native methylotrophs (bacteria and yeasts). Starting with the query protein sequences of *Bacillus methanolicus PB1* Mdh2 and *Pichia angusta* Das, a BLAST search with an identity cut-off at 50% was used with the software CD-HIT to filter out and cluster homologous templates (see Materials and Methods section). This threshold ensured that only functional homologues were identified (Sangar et al., 2007). Some of the 12 prokaryotic Mdh sequences and 17 eukaryotic Das sequences selected in this way belong to genera known to contain methylotrophs, such as *Bacillus, Burkholderia* (Chistoserdova et al., 2009), *Acinetobacter* (Del Rocío Bustillos-Cristales et al., 2017), *Pichia* and *Candida* (Supplementary Figure S2). We added an Mdh from *Bacillus stearothermophilus* and a *P. angusta* methanol oxidase (MOX), since both have been reported to have good affinity for methanol (Kms of 20 mM and 0.4 mM, respectively) (Shleev et al., 2006; Whitaker et al., 2017). Finally we added a Das from *Mycobacterium* with a high affinity for formaldehyde (Km of 1.86 mM) (Ro et al., 1997) and an *E. coli* codon-optimized version of *P. angusta* Das. The Mdh and Das genes were respectively cloned into the low-copy plasmid pSEVA424 and the middle-copy plasmid pSEVA134 (Silva-Rocha et al., 2013). All the selected sequences were assembled in a library of 266 combinations of genes (14 Mdh sequences * 19 Das sequences) and transformed using a robotic platform (Supplementary Figure S3). To ensure that formaldehyde was only assimilated by the pathway introduced into the cell, we used an *E. coli* strain deleted for the first gene of the formaldehyde detoxification pathway (*frmA*)(Figure 1). Since the average optimal growth temperatures of the microbial sources of the Mdh and Das homologues are around 37°C and 30°C, respectively (Supplementary Table S1), we measured enzyme expression at both temperatures. The higher expression obtained at 30°C overnight (Supplementary Figure S4) led us to use this temperature for all subsequent experiments.

To analyze the performance of the 266 different enzyme combinations, methanol incorporation was measured for each pathway variant using dynamic ^13^C-labeling experiments as shown in Supplementary Figure S3. We used the ^13^C-labeling incorporation into the phosphoenolpyruvate (PEP) as a proxy for methanol assimilation since PEP is one of the first multi-carbon products of methanol assimilation (Figure 1). The ^13^C-enrichment of PEP measured for each combination of Mdh and Das is shown in Figure 2A and Supplementary Table S2. ^13^C enrichments of between 1% to 5% were observed for the combinations involving the *das* genes of *Pichia pastoris*, *Verruconis gallopava*, *Scedosporium apiospermum, Rasamsonia emersonii*, *Fonsecaea erecta* and *Kuraishia capsulata*. In comparison, combinations involving the *das* from *P. angusta*, had ^13^C enrichments two to twelve times higher, up to 22% for the codon-optimized version, *P. angusta* (opt), representing an average 2.4-fold increase in ^13^C-isotopic enrichment in PEP compared with the wild type (Supplementary Table S2). These results are consistent with the more stable expression of the codon-optimized version of Das, compared with the wild type version (Supplementary Figure S5).

**Figure 2.**
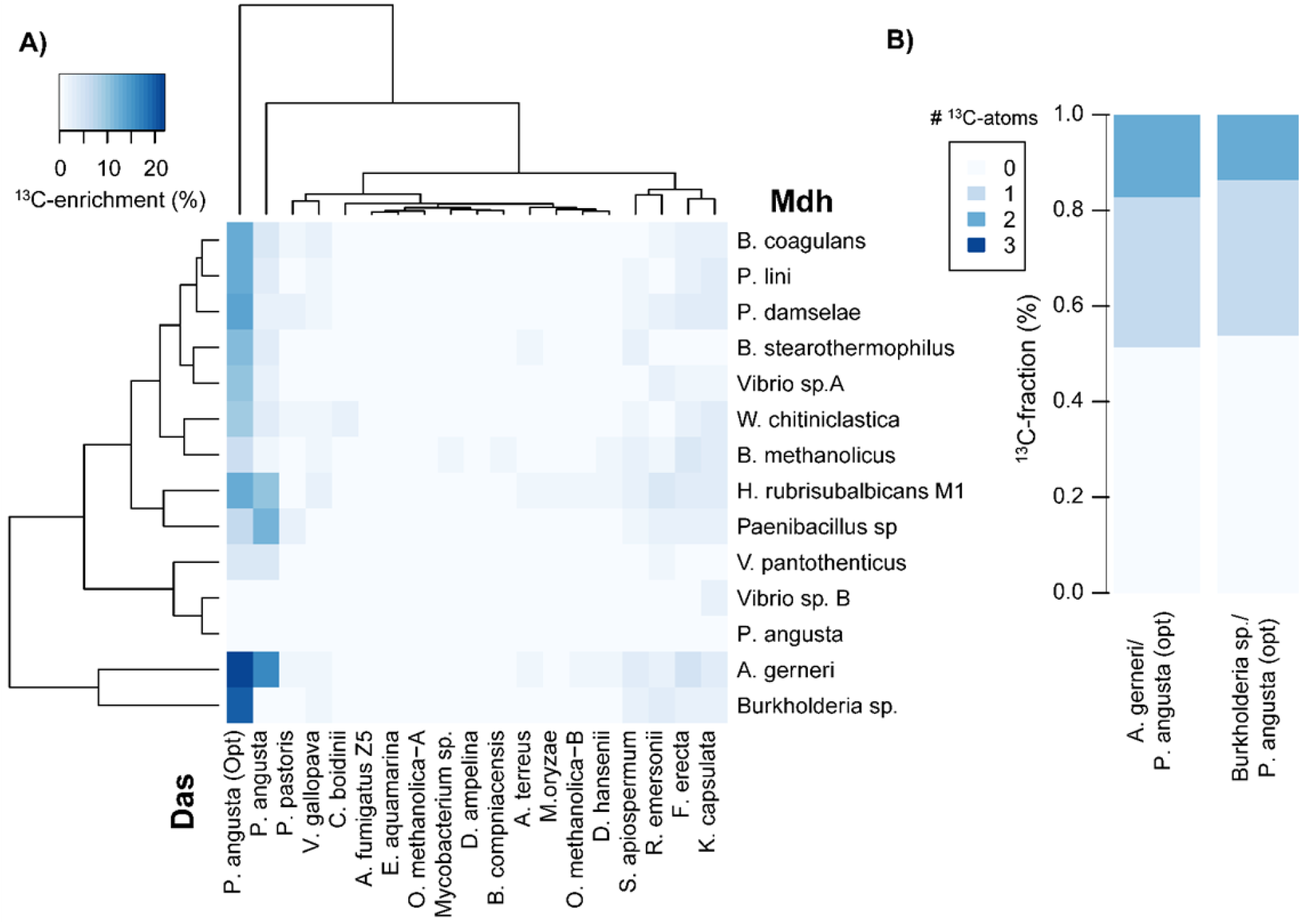
Screening of the different combinations of the hybrid methylotrophic pathway. (A) Heatmap showing the ^13^C-enrichment of phosophoenlypyruvate (PEP) in *E. coli* Δ*frmA* expressing different combinations of Mdh and Das homologues. The ^13^C-enrichment of PEP was measured at the steady state during exponential growth (90 min after cultivation in M9 medium containing 655 mM of ^13^C-methanol). Rows and columns are ordered according to the cluster trees shown on the left and on the top. The Euclidean function was used as distance metric and complete linkage was used as clustering algorithm. (B) Labeling pattern of PEP at 90 min in *E. coli* Δ*frmA* expressing *P. angusta* Das (opt) with either *A. generi* Mdh or *Burkholderia* Mdh.

We then investigated whether methanol assimilation could be increased by optimizing the expression of the Das enzymes which led to low- or non-labeledPEP. The codon-optimized Das genes from *Candida boidinii, P. methanolica A, Aspergillus fumigatus Z5, D. hansenii, R. emersonii* and *K. capsulata* were synthesized and individually co-expressed with *A. gerneri* Mdh. No significant improvement in ^13^C-enrichment was observed compared with the native sequences, except for codon-optimized *R. emersonii* Das, whose PEP labeling was twice as high (4% vs 2% ^13^C-enrichment) (Supplementary Table S3).

Labeling was observed for all these combinations whatever the nature of the Mdh was, suggesting Das compensated for the generally poor kinetic properties of the NAD^+^-dependent Mdh enzymes (Brautaset et al., 2013; Krog et al., 2013) by shifting the equilibrium toward methanol oxidation and subsequent formaldehyde assimilation. However, the level of ^13^C-incorporation was not linked with the expression levels of either Mdh or Das (Figure 2A and Supplementary Figure S5). It is also worth noting that although *B. stearothermophilus* Mdh and *Mycobacterium* Das have favorable *in vitro* and *in vivo* activities (Ro et al., 1997; Whitaker et al., 2017), and were well-expressed in *E. coli* (Supplementary Figure S5), they led to very low ^13^C-enrichement (< 3%) in all tested combinations except those with Das (opt) from *P. angusta* (Figure 2A).

Finally, the highest methanol incorporation was achieved when either *A. gerneri* Mdh or *Burkholderia sp. TSV86* Mdh was expressed in combination with *P. angusta* Das (opt). With these combinations, the ^13^C-enrichements of PEP were respectively 13 and 12 times higher than with the query pathway (*B. methanolicus PB1* Mdh2 / *P. angusta* Das). In particular, the fractions of PEP with one ^13^C atom were 31% and 32% and reached 17% and 13.7% for two ^13^C atoms, respectively (Figure 2B). The incorporation of more than one labeled carbon into PEP demonstrates that the recycling of Xu5P is functional in both combinations.

The higher ^13^C-enrichement obtained for the combination involving *A. gerneri* Mdh and *P. angusta* Das (opt) led us to use these enzymes for subsequent experiments. Although this is the best matching of enzymes, the fact that 100% PEP labeling was not achieved indicates that methanol alone cannot supply all the carbon atoms required for molecular assembly and that pure methylotrophic growth is not yet possible with this pathway.

### 2.3. Characterizing the cellular behavior of the synthetic methylotroph

In order to uncover the specific make-up of the new synthetic methylotrophic *E. coli* strain with regard to methanol utilization, we performed a physiological and transcriptomic analysis of the strain expressing *A. gerneri* Mdh and *P. angusta* Das (opt) grown on xylose with and without additional methanol (Table 1, Figure 3 and raw data in Table S4). For this analysis, both genes were cloned into the pSEVA424 vector as a single operon to avoid the metabolic burdening of the cells with double-antibiotic selection (Silva et al., 2012). Similar levels of ^13^C-methanol incorporation were measured in the single and double plasmid strains after 90 min culture with methanol, but labeling continued to increase in the double plasmid strain up to twice the level observed in the single plasmid strain (Supplementary Figure S6). This can be explained by a decrease in Das levels when the gene is expressed from the low-copy plasmid pSEVA424 (10–15 copies/cell) instead of the middle-copy pSEVA131 plasmid (20–30 copies/cell) (Silva-Rocha et al., 2013).

**Table 1:**
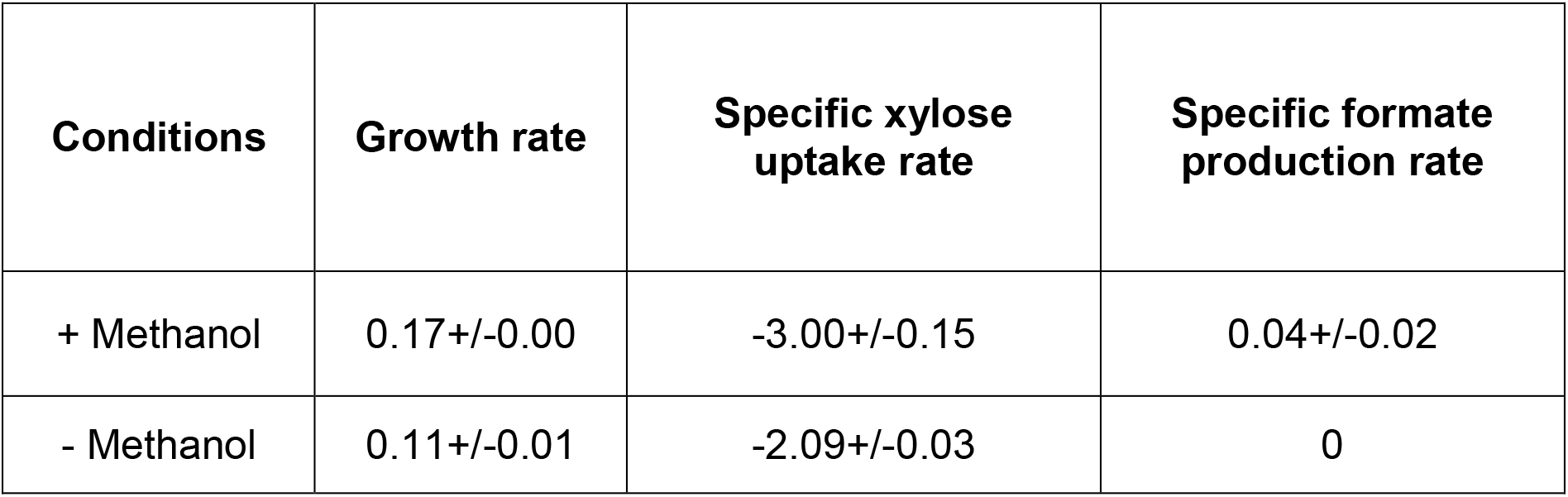
Physiological response of the new synthetic methylotroph to methanol. Growth rate (h^−1^), specific consumption and production rates (mmol/g_DW_/h) of the *E. coli* ΔfrmA_pSEVA424-Mdh-Das(opt) strain during growth in M9 minimal media containing 15 mM xylose without methanol (-Methanol) and supplemented with 150 mM methanol (+ Methanol). Mean and standard deviation of two replicates are given.

**Figure 3.**
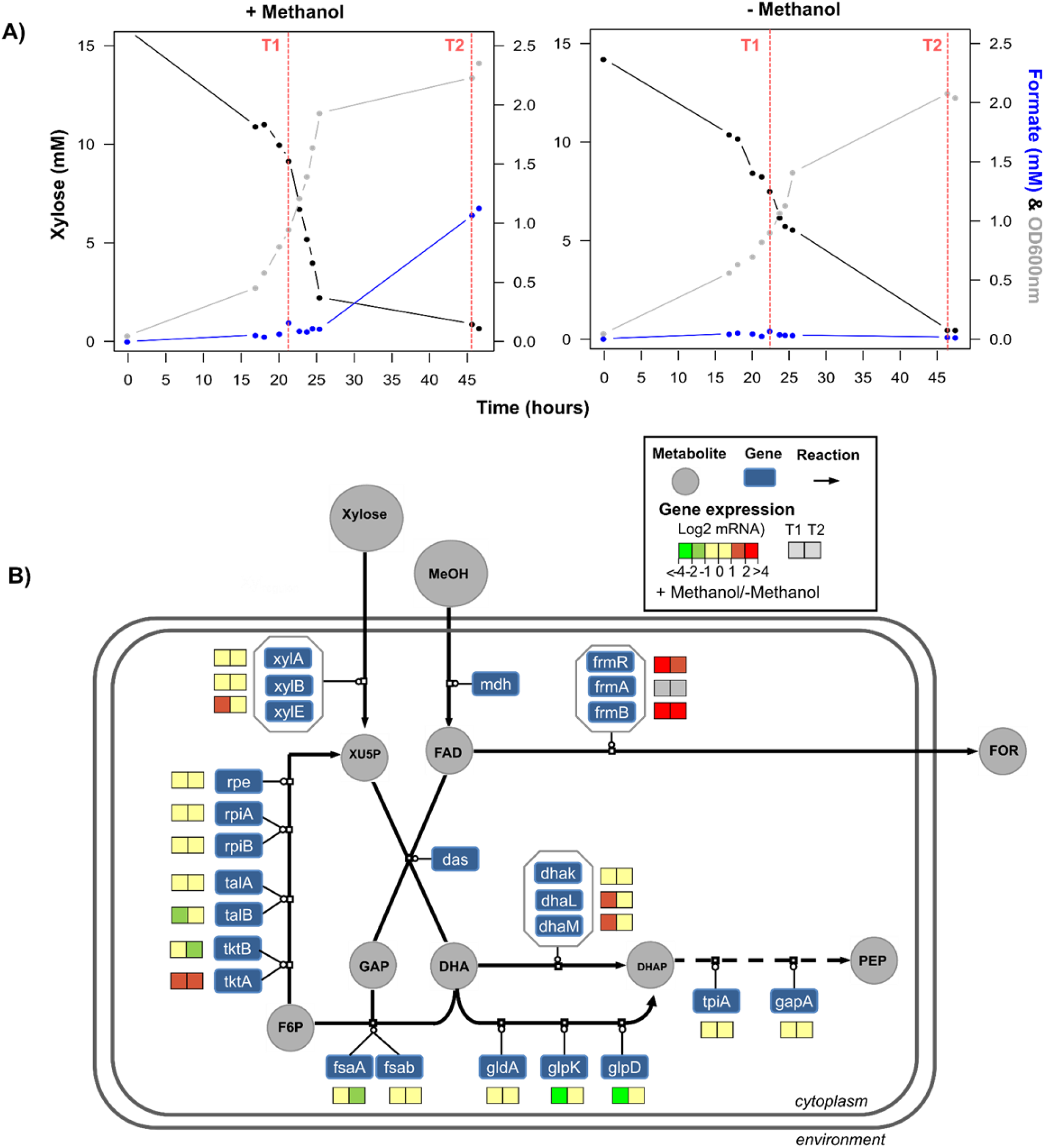
Response of the new synthetic methylotroph to methanol. **A)** Time course analysis of xylose (black), formate (blue) and biomass (gray). Red dotted lines indicates the sampling point (T1 & T2) for transcriptomic analysis. Cells were grown in a minimal synthetic medium containing 15mM xylose with or without 150 mM methanol at 30°C and 250 rpm. One exemplary experiment for each condition is shown (n=2). **B)** Gene expression profiling of the *E. coli* Δ*frmA*_pSEVA424-Mdh-Das(opt). The coloured squares represent the log2-ratios as measure of gene expression fold changes (+Methanol / −Methanol) during exponential growth at OD_600_ = 1 (T1) and when xylose was completely consumed by the cells at OD_600_ = 2 (T2). Methanol dehydrogenase (*mdh*); Dihydroxyacetone synthase (*das*); Glutathione-dependent formaldehyde detoxification operon (*frmRAB);* Dihydroxyacetone kinase operon (*dhaKLM);* Fructose-6-phosphate aldolase isoform A and B (*fsaA, fsaB*); Glycerol dehydrogenase (*gldA*); glycerol kinase (*glpK*); Glycerol-3-phosphate dehydrogenase (*glpD*); Transketolase isoforms A and B (*tktA, tktB);* Transaldolase isoforms A & B (*talA, talB);* Ribose phosphate isomerase isoforms A & B (*rpiA, rpiB);* Ribulose phosphate epimerase (*rpe*); Triose phosphate isomerase (*tpiA*); Glyceraldehyde 3-phosphate dehydrogenase (*gapA*); Xylose isomerase (*xylA);* Xylulokinase (*xylB);* D-xylose/proton symporter (*xylE*); Methanol (MeOH); Formaldehyde (FAD); Xylulose-5-P (XU5P); Glyceraldehyde-3-phosphate (GAP); Dihydroxyacetone (DHA); Phosphoenolpyruvate (PEP); Dihydroxyacetone phosphate (DHAP); Fructose-6-phosphate (F6P); Formate (FOR).

The physiological response to methanol of the synthetic strain expressing the assimilation pathway on one plasmid is given in Table 1. Both the growth rate (+54%) and the specific xylose consumption rate (+45%) were higher in the methanol-supplemented medium than when cells were grown on xylose alone. Formate was only observed in the presence of methanol (Table 1) and its production only started once xylose was depleted (Figure 3A). In contrast, methanol consumption could not be formally assessed since the decrease in concentration occurred at a similar rate as evaporation and fell within the error range of the NMR instrument (4% of the measured value). These results clearly indicate a positive effect of methanol on the rate of xylose uptake, and thus on growth, but also show that formaldehyde was oxidized into formate even though the first step of this pathway had been deleted (i.e. *frmA*).

To characterize the cellular response of the synthetic strain to methanol, a transcriptional analysis was performed at two time points, during exponential growth on xylose (T1) and after xylose exhaustion (T2), with or without methanol supplementation. Specifically, we looked at the expression of the genes involved in methanol metabolism (Figure 3B). In accordance with the observation of formate production only in the presence of methanol (Table 1), the *frmR* and *frmB* genes were strongly up-regulated in the presence of methanol (Figure 3B). While FrmR is a formaldehyde sensor (a proxy for the presence of formaldehyde in the cell), FrmB catalyzes the hydrolysis of the FrmA product S-formylglutathione into formate. Taken together, these data point to the presence of a promiscuous alcohol dehydrogenase that replaces FrmA, but more importantly, confirm the oxidation of methanol into formate. Upregulation of *dhaL* and *dhaM*, which encode the dihydroxyacetone kinase (DAK) pathway, was observed in the presence of methanol, particularly at T1, when xylose was still present (Figure 3B). Because the expression of the *dhaKLM* operon is induced by DHA (Bächler et al., 2005), this confirms the presence of DHA in the cells and thus the co-assimilation of methanol with xylose. However, the genes encoding alternative DHA assimilation routes (i.e. the glycerol (*gldA, glpK* and *glpD*) and the FSA (*fsaA* and *fsaB*) pathways) were not transcriptionally activated or even down-regulated (Figure 3B) at both time points. These results are consistent with the conclusion of Peiro et al. that DHA is mainly assimilated via the dihydroxyacetone kinase (DAK) (Peiro et al., 2019). However, they appear to contradict those of the flux balance analysis that predict that Fsa may be involved in the regeneration of the C1 acceptor. Finally, the gene encoding the transketolase *tktA* was upregulated on methanol at both time points (Figure 3B). This enzyme catalyzes the formation of Xu5P, which plays a key role in the cyclic operation of our synthetic pathway. However, Xu5P is also the entry point of xylose in the metabolism and, interestingly, the presence of methanol improved the expression of *xylE* involved in its transport through the cellular membrane (Figure 3B), particularly during the exponential growth phase. This result corroborates the higher specific xylose uptake rate observed when the synthetic *E. coli* strain was grown in media supplemented with methanol (Table 1).

To assess the global transcriptional changes in response to methanol, we performed Gene Ontology (GO) enrichment analysis. Methanol triggered the up-regulation (≥ 2-fold changes in expression) of respectively 377 and 645 genes and the down-regulation (≤ −2-fold changes) of respectively 177 and 507 genes at T1 and T2, indicating a stronger influence of methanol when xylose was exhausted. The presence of methanol resulted in increased expression of genes related to adhesion and pilus formation (Table 2) during both the exponential and the stationary growth phase. A similar response has been observed in *E. coli* cells grown on DHA (Peiro et al., 2019) and this is in line with the presence of DHA in the cells. Interestingly, among genes related to transcription regulation, positive regulators of the *tdc* operon involved in threonine and serine catabolism were up-regulated (Table 2). This is in accordance with the previous findings that the activation of the threonine degradation pathway is associated with better methanol assimilation (Gonzalez et al., 2018). The presence of methanol had a significant impact on overall cellular organization. The genes encoding ribosomes, translation machinery, amino acid metabolism and other more general processes (Krebs cycle, respiratory chain, fatty acids metabolism, etc.) were down-regulated during the exponential and stationary growth phases (i.e. at T1 and T2) (Table 2).

**Table 2:**
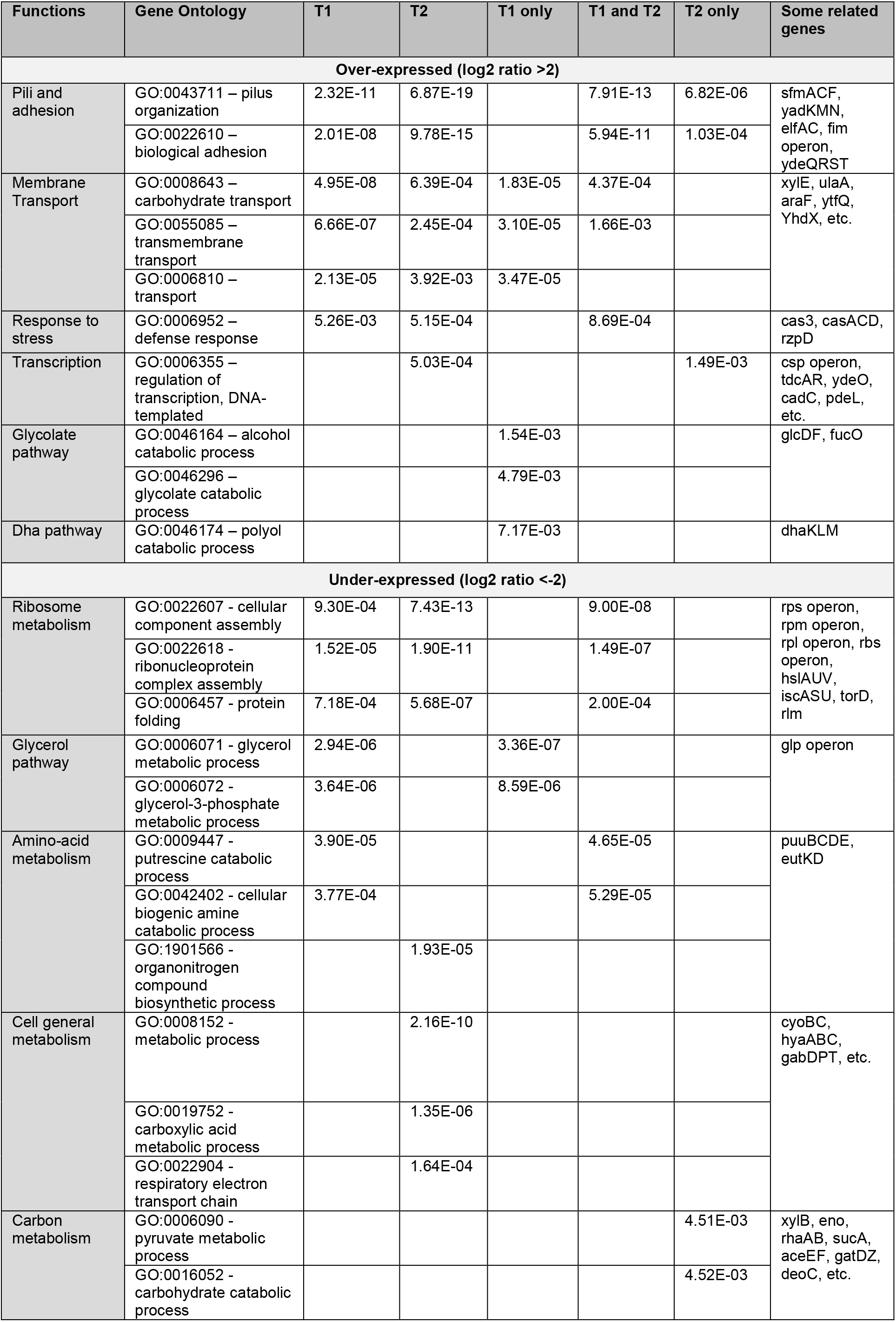
Functional classification of genes with statistically significant increases (log2 ratio >2) and decreases (log2 ratio <-2) in mRNA levels in *E. coli*ΔfrmA_pSEVA424-Mdh-Das(opt) strain during growth in M9 minimal medium containing 15 mM xylose compared to M9-glucose medium supplemented with 150 mM methanol. The categories of orthologous genes (COG) were used for grouping. The indicated values are the p-values attributed to each category on Gene Ontology.

Overall, these data demonstrate that methanol has a negative effect on the cellular machinery by down-regulating cell wall and membrane genes, most probably by chemical perturbation of the outer layers. However, this analysis also shows that methanol can be assimilated by the new synthetic *E. coli* strain and identified genetic engineering targets to limit its dissimilation and improve the cyclic operation of the pathway.

### 2.4. Optimizing the methylotrophic chassis

The above results indicate that genes encoding enzymes for formaldehyde dissimilation (*frmR* and *frmB*), xylulose-5-phosphate recycling (*tktA*) and DHA assimilation (*gldA, glpK, glpD, fsaA* and *fsaB*) are potential targets to boost the assimilation of methanol in our synthetic strain. These genes were thus targeted in the previous strain to engineer a superior methanol assimilation phenotype (Figure 4B). Strain 2 was built by knocking out the entire *frmRAB* operon to avoid drainage of formaldehyde to the detoxification pathway. Strain 3 was built by knocking out the *frmRAB* operon in a Δ*ptsA::kan* mutant in which the *ptsA* gene is replaced by a kanamycin cassette leaving *gldA* and *fsaB* expression under the control of the kanamycin promoter. As a result, *gldA* and *fsaB*, as well as *glpK* and *glpD*, are highly overexpressed compared with the wild-type strain, fully activating both the GLD and FSA pathways (Peiro et al., 2019). In strain 4 finally, *tktA*, a gene encoding a key enzyme in the regeneration of Xu5P, was overexpressed to promote this process.

**Figure 4.**
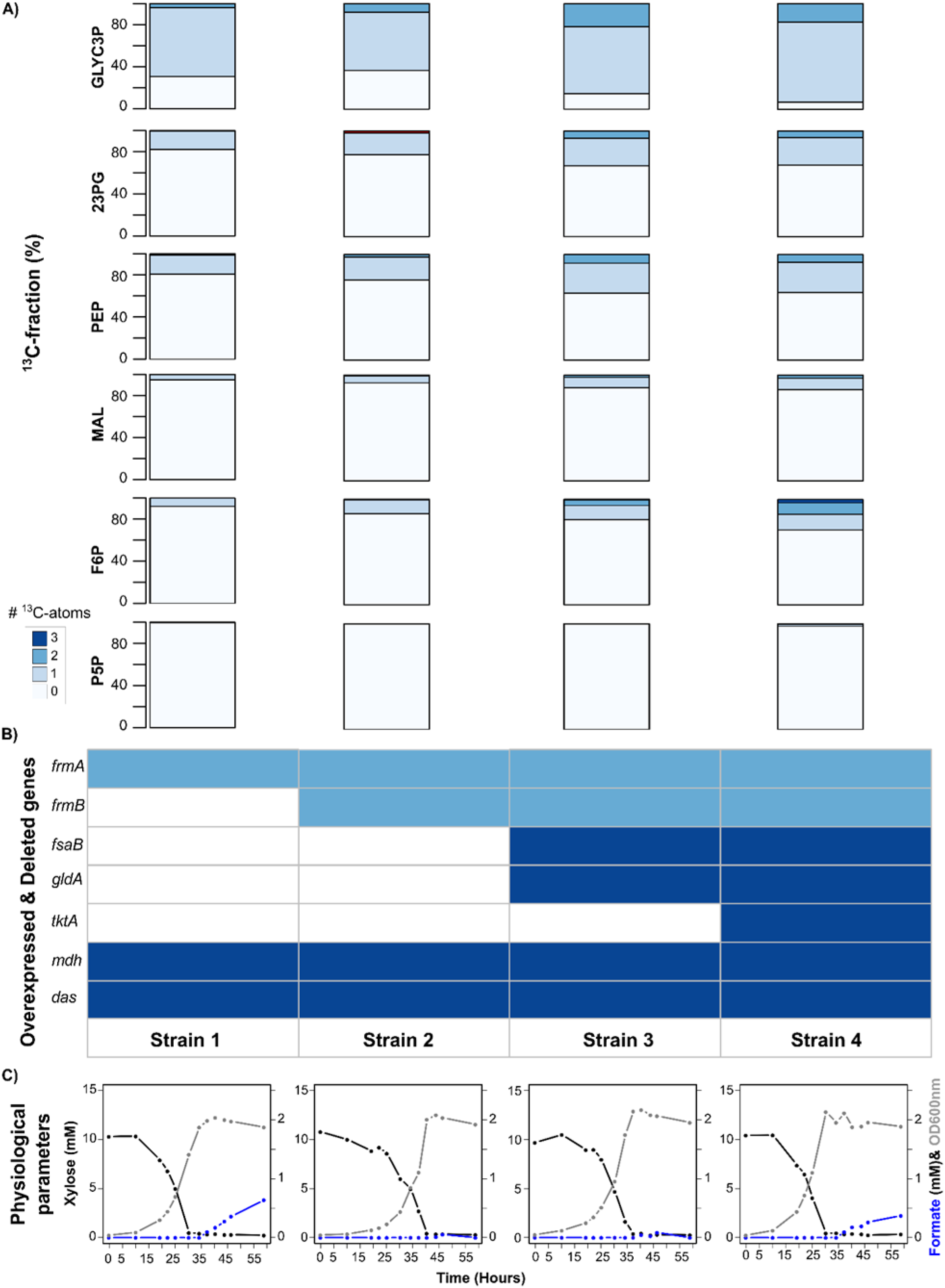
Phenotypic characterization of rationally designed *E. coli* strains. **A)** Labeling patterns of the intracellular metabolites 2 & 3 phosphoglycerate (23PG), fructose-6-phospohate (F6P), glycerol-3-phosphate (GLYC3P), pool of pentoses-5-phosphate (P5P), phosphoenolpyruvate (PEP) and malate (Mal) within the different strains after 90 min of culture in M9 medium with 655 mM ^13^C-methanol. Mean of 3 replicates. **B)** Table of overexpressed (dark blue) and deleted (light blue) genes within the different strains. **C)** Time course analysis of xylose (black), formate (blue) and biomass (gray) within the different strains. Strains were grown in a minimal synthetic medium with 15 mM xylose and 150 mM ^13^C-methanol at 30°C and 250 rpm. One exemplary experiment for each strain is shown (n=2).

To study the impact of these genetic modifications on methanol assimilation, the genealogy of the new rationally designed strains was characterized by following the incorporation of ^13^C-methanol atoms into intracellular and extracellular metabolites (Figure 4 and controls in supplementary Figure S7). Knocking out the *frmRAB* operon (strain 2) resulted in a small increase in ^13^C-methanol incorporation in all the measured intracellular metabolites compared with the starting Δ*frmA* strain 1 (Figure 4A), in line with measurements of the extracellular production of formate (Figure 4C). Upon xylose depletion in the medium, ^13^C-formate production was detected in strain 1 and increased constantly during the stationary phase. In contrast, strain 2 did not produce ^13^C-formate, even after several hours in the stationary phase, confirming that formaldehyde dissimilation is completely prevented in the optimized strain. The most significant improvement in methanol assimilation was observed in strain 3, in which *fsaB* and *gldA* were overexpressed. In this strain, we observed increases of the fractions with one ^13^C atom (M1) in the pool of 2 and 3 phosphoglycerate (23PG, + 8.4%), phosphoenolpyruvate (PEP, +10%) and malate (MAL, +5%) compared with the starting strain (strain 1) (Figure 4A). In line with the activation of the glycerol pathways in strain 3, a large fraction of glycerol-3-phosphate (GLYC3P) with two ^13^C atoms (M2) was measured, an increase of 17.9% compared with strain 1. GLYC3P is an important precursor of membrane constituents and therefore of biomass. In strain 3, all the measured central metabolites had more than one ^13^C atom (Figure 4A), which can only have resulted from recycling of the C1 acceptor, XU5P. This is in accordance with the computational prediction that Fsa plays a key role in the cyclic operation of the synthetic pathway (Figure 1). Strain 4, in which *tktA* was overexpressed, showed even higher ^13^C-methanol incorporation. The fraction carrying two ^13^C atoms was twice as high in F6P compared with strain 3 and a small fraction (3%) containing three ^13^C atoms (M3) was also detected. In addition, a fraction with one ^13^C atom of 2% was measured in the pentose phosphate pool (P5P) containing XU5P. However, ^13^C-formate was once again detected in this strain (Figure 4C).

To confirm that carbon molecules originating from methanol were used in biosynthetic pathways, we also analyzed ^13^C incorporation into proteinogenic amino acids after 48h of cultivation on ^13^C-methanol (Supplementary Figure S8). Low but significant levels of ^13^C were found. In agreement with the labeling observed in the glycolytic and TCA intermediates, labeling was also observed in their derived amino acids i.e. serine (SER, derived from glyceraldehyde-3-phosphate), alanine (ALA, derived from pyruvate), aspartate and glutamate/glutamine (ASP and GLX, derived respectively from oxaloacetate and α-ketoglutarate). As expected from the small amounts of labeled carbon in the P5P pool, no labeling was found in histidine (HIS), which is derived from ribose-5-P. However, some labeling was detected in phenylalanine (PHE), which is derived from erythrose-4-phosphate. The fraction of labeled carbons increased systematically from strain 1 to strain 4 and, more importantly, a fraction of proteinogenic amino acids were found to carry more than one ^13^C atom (Supplementary Figure S8).

## 3. DISCUSSION

Methanol is an attractive feedstock for the production of fuels and chemicals but engineering a C1 fixation pathway into an industrially relevant microorganism, such as *E. coli*, remains challenging. To tackle this problem, this article describes a new computationally designed pathway as an alternative to the well-studied RuMP based pathway (Bennett et al., 2018; Chen et al., 2018; Gonzalez et al., 2018; He et al., 2018; Meyer et al., 2018; Muller et al., 2015; Price et al., 2016; Whitaker et al., 2015; Woolston et al., 2018). This new pathway is a hybrid of naturally occurring cyclic methanol assimilation pathways and consists of a Mdh from *A. gerneri* in combination with a codon-optimized version of *P. angusta* Das. Although the new pathway does not allow the cell to grow on methanol alone, 22% incorporation of methanol carbon was observed in the multi-carbon compound PEP. This is similar to the values measured previously in a synthetic methylotrophic *E. coli* strain expressing cyclic RuMP based-pathways and cultivated under comparable conditions (Supplementary Table S5). Importantly, this article reports the discovery of two novel NAD-dependent alcohol dehydrogenases from Gram-negative, mesophilic, non-methylotrophic organisms (*A. gerneri* and *Burkholderia* sp.) with significant *in vivo* affinity for methanol. Representatives of the Burkholderia order have recently been recognized as true facultative methylotrophs (Chistoserdova et al., 2009) and one NAD-dependent Mdh from this order has the highest *in vitro* affinity for methanol reported to date (Woolston et al., 2018; Wu et al., 2016; Yu and Liao, 2018). In our setting, the two novel Mdhs performed better *in vivo* than the Mdhs from *B. methanolicus* and *B. stearothermophilus* which were used previously to implement methylotrophy in *E. coli* (Bennett et al., 2018; Chen et al., 2018; Gonzalez et al., 2018; Kim et al., 2020; Meyer et al., 2018; Muller et al., 2015; Whitaker et al., 2017). However, the efficiency of the pathway was mostly improved by using a codon-optimized version of Das, indicating that this enzyme is very likely rate-limiting for methanol assimilation. Since Das was not overexpressed as much as Mdh was (Supplementary Figure S4), further increasing its expression should also increase methanol assimilation.

Our iterative process of strain analysis and engineering combining omics and modelling approaches was decisive in the selection of strategic genetic targets to maximize methanol assimilation. The final optimized strain incorporated 1.5 to 5.9 times more methanol — as measured by ^13^C-enrichment and depending on the metabolite — than did the starting strain. A maximum ^13^C-enrichment of 37% was achieved in GLYC3P. In addition, the increase in the number of labeled carbons per molecule for most metabolites shows that cyclic operation of the synthetic pathway was improved in the final strain. Finally, the presence of labeling in biomass constituents showed that carbon molecules originating from methanol were not only assimilated into the central metabolism but also used in biosynthetic pathways. This is evidence of true methanol metabolism and confirms the establishment of methylotrophy in this *E. coli* strain. In the optimized strain, the most significant improvement was achieved by activating alternative DHA assimilatory pathways. This is consistent with a previous study demonstrating that the specific DHA uptake rate in a similar engineered strain was increased by 60% (Peiro et al., 2019). We further improved methanol assimilation in the synthetic strain by overexpressing a transketolase and, therefore, improving the recycling of the C1 acceptor. This is in agreement with the conclusion of a previous study that expressing the non-oxidative pentose phosphate pathway (PPP) from *B. methanolicus* improves methanol assimilation in a synthetic *E. coli* methylotroph (Bennett et al., 2018).

Finally, we also observed that methanol improved the growth of our synthetic strain on xylose by up-regulating the genes involved in xylose transport through the cellular membrane. Up-regulation of genes encoding transmembrane transporters in the presence of methanol has also been observed in *S. cerevisiae* (Espinosa et al., 2019). The chemical properties of methanol are known to modify the physical properties of cell membranes, such as their fluidity (Joo et al., 2012). These changes can be perceived by the cells and trigger the expression of genes that are involved in the acclimation of cells to new conditions (Los and Murata, 2004). However, we also observed that all the basic processes necessary for cell growth were negatively affected by the presence of methanol. The fact that so many cellular systems were down-regulated is a further indication that methanol alters the physical properties of cell membranes, triggering the associated gene expression response, the membrane being the first point where toxic effects can occur (Denich et al., 2003).

In this work, we successfully created an *E. coli* strain able to efficiently assimilate methanol through a brand new synthetic metabolic pathway. However, there is still room for optimization and our results suggest that the overall metabolic capacity for methanol can be improved in several ways. For example, one could (i) improve the expression of Das, (ii) block all the dissimilatory pathways, (iii) improve the recycling of the C1 acceptor, and (iv) coordinate the catabolic pathway with the overall cellular infrastructure by engineering methanol-sensitive elements to improve the global response to the substrate (Rohlhill et al., 2017) or by directed evolution (Chen et al., 2018; He et al., 2018; Meyer et al., 2018). However, a recent study demonstrating the slow growth (doubling time of 54 h) on a mixture of methanol and CO2 of an *E. coli* strain expressing a linear methanol assimilation pathway (Kim et al., 2020) raises questions about the relevance of establishing methylotrophy in *E. coli* using cyclic pathways. Arguments in favor of pursuing the quest for growth on pure methanol using cyclic pathways are (i) the independence of such pathways from other carbon sources, and (ii) a recent study reporting an *E. coli* strain expressing an autotrophic cycle capable of producing all its biomass carbon from CO2 (Gleizer et al., 2019).

## 4. Materials and Methods

### 4.1 Reagents

All chemicals were purchased from Sigma-Aldrich (St. Louis, MO, USA) unless noted otherwise. Unlabeled methanol (≥ 99.9%, LC-MS grade) was purchased from Honeywell (Muskegon, MI, USA). Isotopically labeled ^13^C-methanol (99% ^13^C) was purchased from Eurisotop (Saint-Aubin, France). Phusion^®^ DNA polymerase and restriction enzymes were purchased from New England Biolabs Inc. (Beverly, MA, USA).

### 4.2 Bacterial strains and culture media

All the strains, plasmids, primers and synthetic gene constructs used in this study are listed in Supplementary Table S6. *E. coli* DH5α was used for plasmid construction and propagation whereas *E. coli* BW25113 was used for methanol assimilation. *E. coli* BW25113Δfrma::*neo* was obtained from the Keio collection and the Flp recognition target (FRT)-flanked kanamycin cassette was removed using Flp recombinase from pCP20 plasmid (Cherepanov and Wackernagel, 1995). After recombination, loss of pCP20 was confirmed by re-streaking on ampicillin, and removal of the resistance cassette was confirmed by polymerase chain reaction (PCR). For operon construction, *A. gerneri* Mdh and *P. angusta* Das genes, containing RBS and a 6xHis tag, were amplified from the pSEVA plasmids using primers P91&P92 and P83&P93, respectively. The two fragments, designed to overlap by 35 bp, were joined by overlapping PCR. The complete Mdh-Das operon was subsequently cloned into the pSEVA424 vector using primers P102&P103 and the In-Fusion^®^ HD kit (Takara Bio, Otsu, Japan). The λ red recombination method (Datsenko and Wanner, 2000) was used to generate knockout strains ΔptsA (primers P129&P130) and ΔfrmRAB (primers P145&P146). The introduced antibiotic resistance cassettes were removed using the FRT/FLP recombination system (Cherepanov and Wackernagel, 1995). All constructs were subsequently verified by colony PCR and sequencing (GATC, Konstanz, Germany).

All *E. coli* strains harboring plasmids were propagated in Luria-Bertani (LB) medium or M9 minimal medium containing the appropriate antibiotics. The composition of the M9 minimal medium was as follows (in g·L^−1^): 18 Na_2_HPO_4_, 3.13 KH_2_PO_4_, 0.53 NaCl, 2.11 NH_4_Cl, 0.49 MgSO_4_·7H_2_O, 0.00438 CaCl_2_·2H_2_O, 0.1 thiamine hydrochloride, trace elements (mg L-1) 15 Na_2_EDTA·2H_2_O, 4.5 ZnSO_4_·7H_2_O, 0.3 CoCl_2_·6H_2_O, 1 MnCl_2_, 1 H_3_BO_3_, 0.4 Na_2_MoO_4_·2H_2_O, 3 FeSO_4_·7H_2_O, 0.3 CuSO_4_·5H_2_O. The antibiotics were added when necessary in the following concentrations: ampicillin (Amp, 100 μg/ml), kanamycin (Kan, 50 μg/ml), streptomycin (Strp, 50 μg/ml). The optical density at 600 nm (OD600) was measured using a GENESYS 6™ spectrophotometer (Thermo Scientific).

### 4.3 In silico design of the synthetic pathway

The synthetic pathway for methanol assimilation was designed using the software FindPath (Vieira et al., 2014). The workflow starts with the creation of a substrate-associated reaction database based on the literature and available metabolic databases. This database consists of reactions involving the target molecule (in our case, methanol). The database is then converted into a model that is subsequently used to compute elementary flux modes (EFMs), i.e., all the possible flux distributions in a metabolic network under steady state conditions. Among these EFMs, the best pathways are selected and ranked according to their efficiency. Finally, the best module combinations for efficient methanol conversion were identified. In our case, the methanol database encompassed more than 100 reactions steps and 100 metabolic compounds involved in methanol metabolism. For each reaction, the genes, reaction, EC number, KEGG name, localization, and reversibility were reported. Finally, the model was built by bringing together all the reactions along with the transporters and cofactor recycling, i.e. 47 reactions and 114 metabolites. Using this model, 10000 EFMs were generated of which 85 allowed the conversion of methanol into *E. coli* metabolites. From these, 20 efficient EFMs were selected, i.e. those involving a small number of reactions and with low cofactor consumption (ATP, NAD(P)H). The hypothesis was that their introduction into the host would require little genetic effort (the number of genes being correlated with the number of reactions) and would have little or no effect on the host’s energy and redox machinery. The reactions composing the 20 EFMs were implemented in a genome scale model *E. coli* (iAF1620). Finally, the biomass yields on methanol of each of the 20 EFMs were simulated using *in-silico* flux balance analysis (FBA),.

### 4.4 Library generation by combinatorial assembly

A BLAST search against UniRef50 (Suzek et al., 2014) using *B. methanolicus* PB1 Mdh2 (UniProt ID: I3DVX6) and *P. angusta* Das (UniProt ID: P06834) as query sequences returned two clusters with 177 and 230 members, respectively. The sequence clustering tool H-CD-HIT (Huang et al., 2010) was used to hierarchically merge similar sequences at varying levels of sequence identity. Proteins were first clustered at a high identity (90%) before the non-redundant sequences were further clustered at a low identity (80% and eventually 70%). Among the representatives of the different clusters, we selected 12 putative Mdh variants and 17 putative Das variants from aerobic and mesophilic microorganisms. The corresponding Mdh genes, as well as the *Bacillus stearothermophilus* Mdh (Dowds et al., 1988) and *Pichia angusta* Mox genes (Shleev et al., 2006), were cloned in the expression vector pSEVA424 (Silva-Rocha et al., 2013) between restriction sites AvrII and NotI. The selected Das genes, plus the *Mycobacterium* Das gene (Ro et al., 1997) and an *E. coli* codon-optimized version of P06834, were cloned in the expression vector pSEVA134 (Silva-Rocha et al., 2013) between restriction sites AvrII and SpeI. All the constructs were synthesized and cloned by BaseClear (Leiden, The Netherlands). The same ribosome binding site (RBS) (<u>AGGAGG</u>AAAAACAT) and 6xHis tag was used for all the genes. The two gene libraries were co-transformed in the BW25113ΔfrmA::frt strain using the rubidium chloride method (Green and Rogers, 2013) and plated on LB-Amp-Strp plates (Supplementary Figure S3).

### 4.5 Dynamic ^13^C-labeling incorporation

To study the incorporation of ^13^C-methanol into intracellular metabolites and proteinogenic amino acids, cells were first cultured in M9 minimal medium in the presence of 15 mM xylose, antibiotics and 0.1 mM IPTG, in 96-deep-well plates, at 30°C and 220 rpm until exponential phase (OD600 = 0.5-1). The cells were then centrifuged at 4400g for 3 min and resuspended in M9 minimal medium with reduced (five times less) phosphate and sulfate, IPTG, antibiotics, and ^13^C-methanol (655 mM). The methanol concentration was chosen to be sufficiently above the Km of Mdh.

After 90 and 180 min incubation at 30°C and 220 rpm, intracellular metabolites were sampled as follows: 120 μL of culture was taken and mixed with 1 mL of cold (−20°C) acetonitrile:methanol:water:formic acid (40:40:20:0.1) extraction solution. The samples were vacuum-dried overnight. The next morning, dried metabolites were resuspended in 120 μL water, centrifuged at 16,000 × g for 2 min, and injected into the LC-MS. Central metabolites were separated on a Dionex™ IonPac AS11-HC anion-exchange column (250 × 2 mm) equipped with an AG11 guard column (50 × 2 mm) with KOH as the mobile phase using a Dionex™ ICS-5000+ Reagent-Free™ HPIC™ system (Thermo Fisher Scientific™, Sunnyvale, CA, USA). Separation of PEP shown in Figure 2 was carried out with a flow rate set at 0.38 ml/min and the following elution gradient: 0 min, 0.5 mM; 1 min, 0.5 mM; 9.5 min, 4.1 mM; 14.6 min, 4.1 mM; 24 min, 9.65 mM; 31.1 min, 100 mM and 43 min, 100 mM. For separation of central metabolites shown in Figure 4, the elution gradient was as follows: 0 min, 7 mM; 1 min, 7 mM; 9.5 min, 15 mM; 20 min, 15 mM; 30 min, 45 mM; 33 min, 70 mM; 33.1 min, 100 mM; 42 min, 100mM; 42.5 min, 7 mM and 50 min, 7 mM. Metabolites were detected using a Thermo Scientific™ LTQ Orbitrap Velos™ mass spectrometer in negative electrospray ionization mode. The spray voltage was 2.7 kV, the capillary and desolvatation temperatures were 350°C, and the maximum injection time was 50 msec. The spectrometer was operated in full-scan mode at a resolution of 60,000 (400 m/z).

After 48 h of incubation at 30°C and 220 rpm, proteinogenic amino acids were sampled as follows: the plates were centrifuged at 4400g for 3 min and the supernatant was removed. To release protein-bound amino acids from cellular proteins, the cell pellets collected were hydrolyzed for 15 h with 6N HCl at 100°C. HCl was evaporated at low pressure (20 mbar, room temperature). Biomass hydrolysates were washed twice in water using the same evaporation method. The dried hydrolysates were resuspended in 200 μL water and centrifuged. A 10-fold dilution was prepared, and samples were analyzed by LC-HRMS. Proteinogenic amino acids were separated on a Supelco™ HS F5 DISCOVERY column (150 × 2.1 mm; 5 μm) equipped with a SUPELGUARD KIT HS F5 guard column (20 × 2.1 mm; 5 μm) with 0.1% formic acid (solvent A) and 0.1% acetonitrile/formic acid (solvent B) as the mobile phase using a UHPLC Vanquish system (Thermo Fisher Scientific™, Sunnyvale, CA, USA). The flow rate was set to 0.25 ml/min and the elution gradient was (% B): 0 min at 2%, 2 min at 2%, 10 min at 5%, 15 min at 35%, 20 min at 100%, 24 min at 100%, 24,1 min at 2% and 30 min at 100%. Metabolites were detected using a Thermo Scientific™ Orbitrap Q-Exactive+™ mass spectrometer in positive electrospray ionization mode, with a spray voltage of 5 kV, and capillary and desolvatation temperatures of 250°C. The spectrometer was operated in full-scan mode at a resolution of 60,000 (400 m/z).

^13^C-carbon isotopologue distributions were identified by matching masses from the mass spectra (mass tolerance of 5 ppm) and retention times using the software TraceFinder (v. 4.1). The peaks of different isotopologues were integrated and corrected for the natural abundance and isotopic purity of the tracer using the software IsoCor (Millard et al., 2019). Levels of ^13^C-isotopic enrichment were then determined as follows: ^13^C-enrichement (%) = sum(Mi*i)/n, where n is the number of carbon atoms for the measured fragment and Mi is the corrected abundance of the mass isotopologue.

### 4.6 Supernatant analysis

Metabolite utilization and the production of the synthetic methylotroph were analyzed by quantitative 1D ^1^H-NMR at 280 K using a zgpr30 sequence with water pre-saturation prior to acquisition on an Avance III 500 MHz spectrometer (Bruker, Rheinstetten, Germany) equipped with a 5 mm QPCI cryogenic probe head. The parameters were as follows: 286°K, 128K points, 8 s relaxation time, 2 dummy scans, 32 scans. Free induction decays (FIDs) were converted into frequency domain spectra by Fourier transform. All spectra were processed using the software TopSpin (v. 3.5). Phases were adjusted manually, baselines were adjusted automatically, and the spectra were aligned and quantified using 3-trimethylsilylpropionic-2,2,3,3-d4 acid sodium salt (TSP-d4, 1 mM) as a chemical shift and concentration standard. The concentrations of the different metabolites (xylose, methanol, formate, and acetate) were calculated with the following equation: concentration = integrated peak area*TSP concentration*dilution of the sample/number of protons in the molecule. For xylose, only the peaks corresponding to the anomeric protons were integrated.

### 4.7 Transcriptomic analysis

Cells were grown in flasks of M9 minimal media containing 15 mM xylose with or without 150 mM MeOH. At T1 (OD600 = 1, exponential phase) and T2 (OD600 = 2, stationary phase), 4 mL of each culture was centrifuged for 90 s at 14000 rpm before discarding the supernatant and immediately freezing the pellets in liquid nitrogen. Total RNA was extracted according to the Qiagen RNAeasy MiniKit procedure and quantified using a Nanodrop^®^ spectrophotometer. Double-stranded complementary DNA (cDNA) synthesis and array processing were performed using the Agilent Technologies One-Color Microarray-Based Gene Expression Analysis protocol. The images were analyzed with the software DEVA (v. 1.2.1). All array procedures were performed using the GeT-Biopuces platform (http://get.genotoul.fr/). For each data set, corresponding to time point T1 or T2, the log2 intensities obtained in the presence of methanol were divided by the log2 intensities obtained without methanol. These ratios were then normalized by the log median intensity. Genes whose expression level differed by a factor of 2 or more between the two conditions were selected for further analysis. Gene ontology analyses were performed using Ecocyc (https://ecocyc.org/). Gene expression data have been deposited in the ArrayExpress database at EMBL-EBI (www.ebi.ac.uk/arrayexpress) under accession number E-MTAB-8909.

### 4.8 In silico analysis of methanol metabolism

We used the functions “flux balance analysis” (FBA) and “flux variability analysis” of the R environment (R Development Core Team, 2009; Team, 2015) Sybil Package (Gelius-Dietrich et al., 2013) and the genome scale model of *E. coli* ij01366 (Orth et al., 2011) amended with the heterologous reactions catalyzed by Mdh, Das and Glpx, and their associated metabolites to simulate the growth and fate of methanol. The objective function was the growth rate whereas the model was constrained using the methanol uptake rate measured experimentally for the wild-type methylotroph *Methylobacter extorquens* (15 mmol/gW/h).

### 4.9 Growth and methanol consumption calculations

Specific growth rates, uptake rates and production rates were determined using PhysioFit, provided open source at https://github.com/MetaSys-LISBP/PhysioFit. A conversion factor of 0.37 g dry weight/OD600 was used.

## 5. Acknowledgment

The authors thank MetaToul-MetaboHUB (Metabolomics & Fluxomics Facitilies, Toulouse, France, www.metatoul.fr, ANR-11-INBS-0010, www.metabohubmetabohub.fr) and its staff members for technical support and access to MS and NMR facilities. This work was funded by the ANR (ANR-16-CE20-0018-01 & ANR-17-COBI-0003-05) and TWB (https://www.toulouse-white-biotechnology.com/). The authors also thank H. Cordier and O. Galy from TWB for technical support and access to robotic platform, and the following students, L. Balcells, M. Beaudor, C. Brodeau, M. Cayet, T. Chaillet, C. Ledoux and E. Sylvander for their help producing transcriptomic data and initiating the data analysis.

## 6. Author contribution

A. De Simone, C.M. Vicente and C. Peiro built the strains and performed the physiological experiments. L. Gales and F. Bellvert performed the MS analysis. B. Enjalbert performed the transcriptomic analysis. S. Heux designed the study and wrote the paper with the help of all the co-authors. The authors declare that they have no conflicts of interest.

## 8. Supplementary Materials

As noted in the text, supplementary materials are available in the online version of this paper. Supplementary Figures contains Figures S1 to S8.Figure S1 shows and overview of the recycling of the C1 acceptor Xu5P and its operation. Figure S2 shows the unrooted phylogenetic trees of selected Mdh and Das homologues. Figure S3 shows the overall scheme of the combinatorial assembly and screening of the synthetic pathway. Figure S4 shows the expression analysis of *B. methanolicus* Mdh and *P. angusta* Das in different conditions. Figure S5 shows the western Blot analysis of expression of Mdh and Das homologues.Figure S6 shows the ^13^C-Methanol assimilation in the methylotrophic *E. coli* Δ*frmA* expressing the synthetic pathway from one or two vectors. Figure S7 shows the ^13^C-Methanol assimilation into central metabolism intermediates in the control strains of the genealogy of methylotrophic *E. coli*. Figure S8 shows ^13^C-Methanol assimilation into proteinogenic amino acids in the genealogy of methylotrophic *E. coli*.

Supplementary Tables contains Tables S1 to S6. Table S1 is the list of selected Mdh and Das homologues and the the associated optimal growth temperature range of the source organisms. Table S2 contains the mean isotopic enrichment of PEP in % using the combinatorial library. Table S3 contains the mean isotopic enrichment of PEP using different Das enzymes with codon-optimized sequences in combination with MDH from *A. gerneri*. Table S4 contains the differential gene expression in the *E. coli* Δ*frmA*_pSEVA424-Mdh-Das(opt) strain during growth in M9 minimal media containing 15 mM xylose without methanol and supplemented with 150 mM methanol during exponential growth at OD600nm =1 (T1) and when xylose was completely consumed by the cells at OD600nm=2 (T2). Table S5 contains an overview of the ^13^C-enrichement obtained in different synthetic methylotrophic strains from previous studies. Table S6 is the list of stains and plasmids used in this study.

