## Supplementary material for "Mixing and matching methylotrophic enzymes to design a novel methanol utilization pathway in *E. coli*": Supplmentary Figures

### Mixing and matching methylotrophic enzymes to design a novel "hybrid" metabolic pathway for methanol assimilation in *E. coli*

De Simone A.[1]; Vicente C.M.[1]; Peiro C.[1]; Gales L.[1,2], Bellvert F.[1,2]; Enjalbert B.[1]; Heux S.[1*]

[1]TBI, Université de Toulouse, CNRS, INRAE, INSA, Toulouse, France
[2]MetaboHUB-MetaToul, National Infrastructure of Metabolomics and Fluxomics, Toulouse, 31077, France

#### Supplementary Figures S1 to S8.

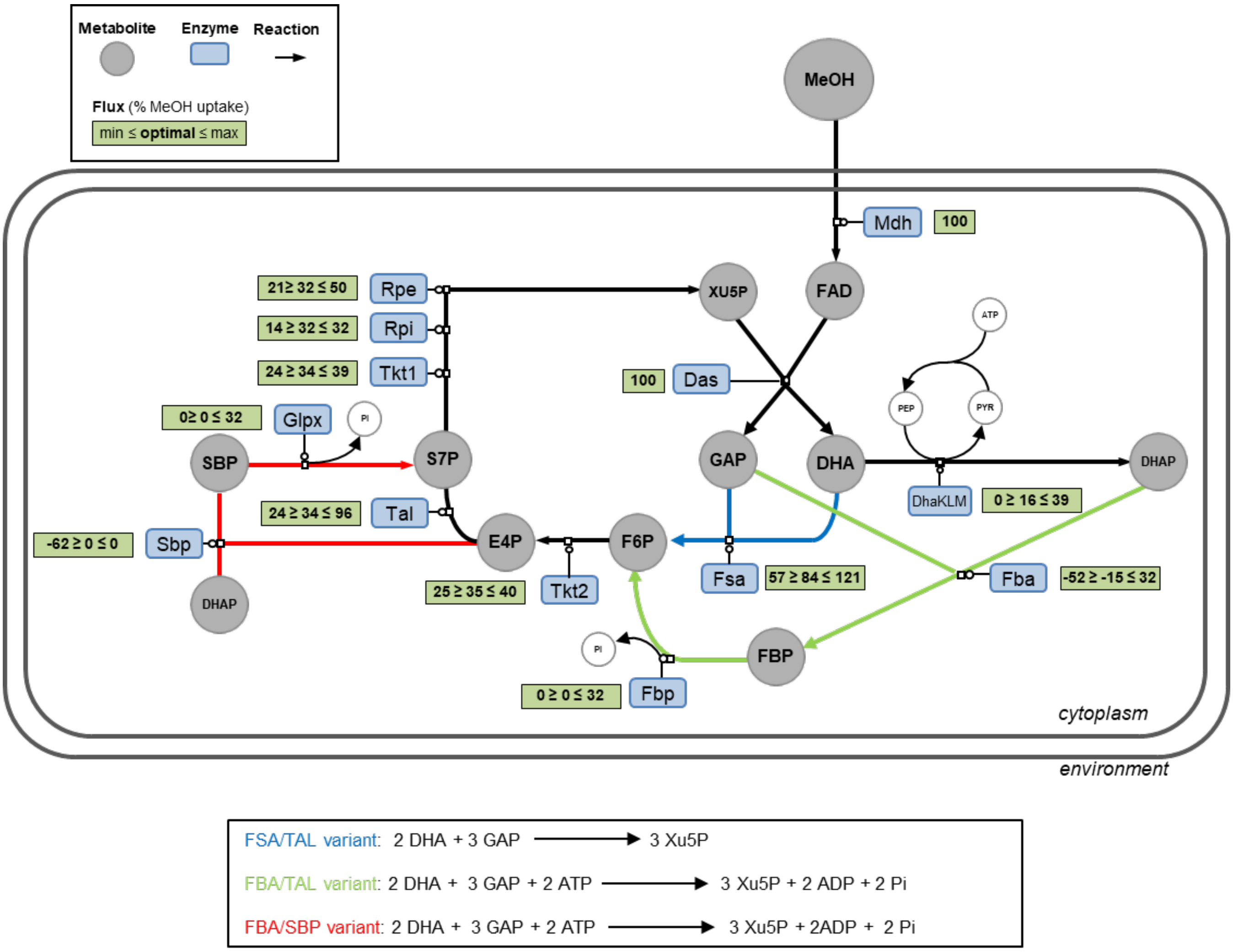


**Supplementary Figure S1. Overview of the recycling of the C1 acceptor Xu5P and its operation.** Recycling can occur through three possible sets of rearrangement reactions which differ in their ATP requirements: the *in silico* predicted FSA/TAL variant (blue lines), and the alternative FBA/TAL (green lines) and FBA/SBP variants (red lines). Green rectangles give the optimal and the ranges of simulated fluxes obtained using flux balance analysis and flux variability analysis, respectively, when growth rate is constrained to 90% of the optimal value. Flux values are given in % relative to a MeOH uptake rate of 15 mmol/gDW/h as defined in Peyraud et al., BMC Syst Biol. 2011.

Dihydroxyacetone kinase (DhaK, DhaL and DhaM); Glycerol dehydrogenase (GldA); Glycerol-3-phosphate dehydrogenase (GlpD); Glycerol kinase (GlpK); Fructose-6-phosphate aldolase (FsaA and FsaB); Triose phosphate isomerase (Tpi); Glyceraldehyde-3-phosphate dehydrogenase (Gapd); Ribulose 5-phosphate 3-epimerase (Rpe); Transketolase (Tkt1 & Tkt2); Transaldolase (Tal); Ribose-5-phosphate isomerase (Rpi); Fructose-bisphosphate aldolase (Fba); Fructose 1,6-bisphosphatase (Fbp); Sedoheptulose bisphosphatase (Sbp). Methanol (MeOH); Formaldehyde (FAD); Xylulose-5-P (XU5P); Glyceraldehyde-3-phosphate (GAP); Dihydroxyacetone (DHA); Phosphoenolpyruvate (PEP); Dihydroxyacetone phosphate (DHAP); Fructose-6-phosphate (F6P); Fructose bisphosphate (FBP); Erythrose 4-phosphate (E4P); Sedoheptulose 1,7-phosphate (SBP); sedoheptulose-7-phosphate (S7P).

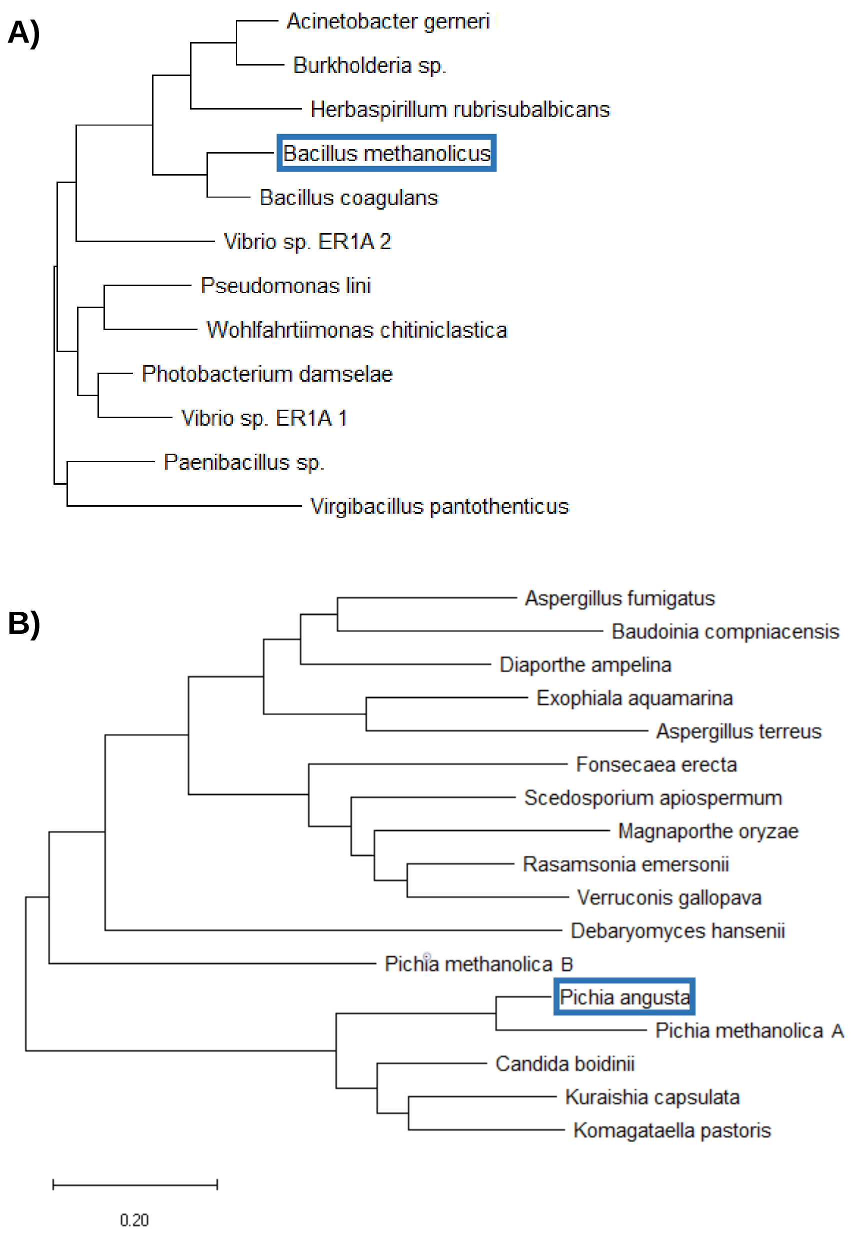


**Supplementary Figure S2. Unrooted phylogenetic trees of selected Mdh (A) and Das (B) homologues.** The evolutionary history was inferred by using the Maximum Likelihood method and Le_Gascuel_2008 model [1]. The unrooted tree with the highest log likelihood (-6124.84) is shown. The tree is drawn to scale, with branch lengths measured in the number of substitutions per site. Evolutionary analyses were conducted in MEGA X [2]. The query sequences are highlighted in blue. The sequences of *A. gerneri* Mdh and *Burkholderia* Mdh share 77.2% identity.

1. Le S.Q. and Gascuel O. (2008). An Improved General Amino Acid Replacement Matrix. Mol Biol Evol 25(7):1307-1320.
2. Kumar S., Stecher G., Li M., Knyaz C., and Tamura K. (2018). MEGA X: Molecular Evolutionary Genetics Analysis across computing platforms. Molecular Biology and Evolution 35:1547-1549.

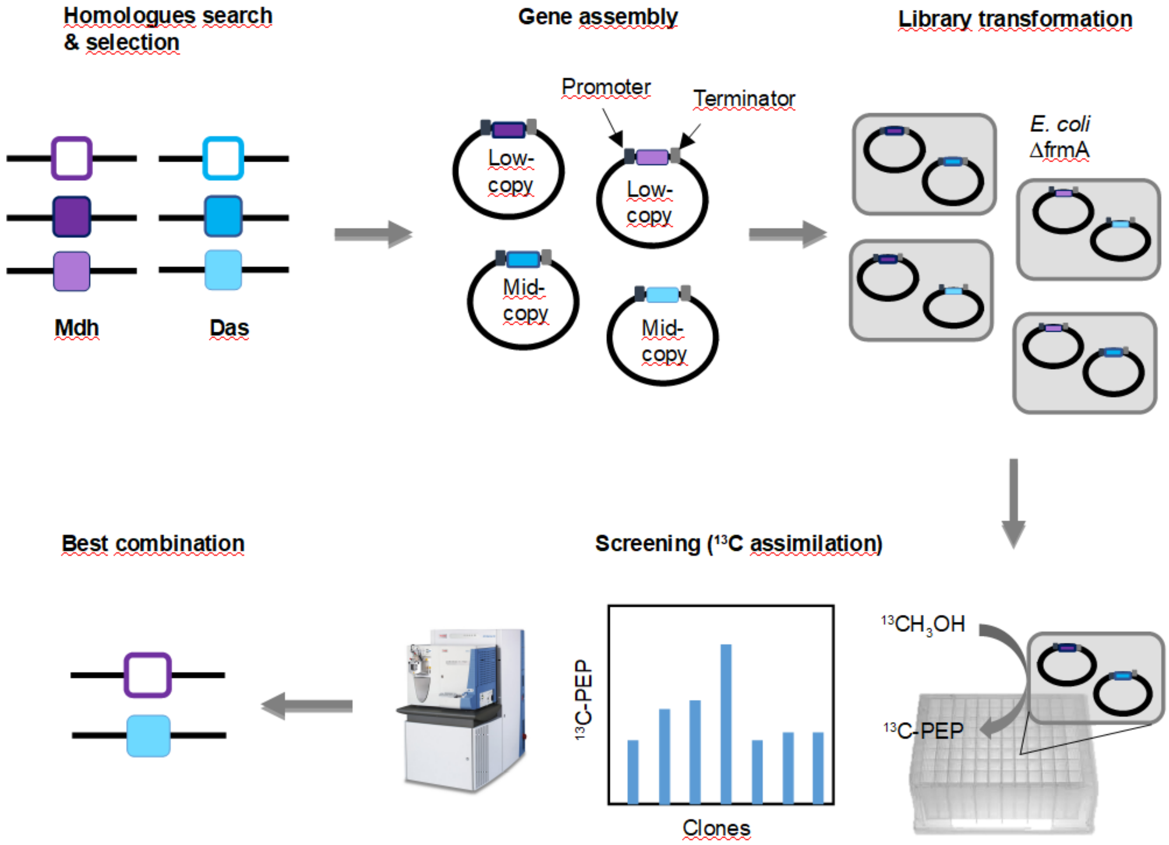


**Supplementary Figure S3. Overall scheme of the combinatorial assembly and screening of the synthetic pathway.** Genes encoding Mdh and Das homologues are synthesized and cloned into two different plasmids between a promoter and terminator. A library is created by mixing the two arrays of plasmids and co-transforming each two-gene combination into the selected strain. The transformed strains are individually cultured in presence of $^{13}$C-methanol and screened by measuring $^{13}$C incorporation levels into PEP, a direct metabolite of the synthetic pathway. The best combination is identified based on the highest $^{13}$C-incorporation level.

A)

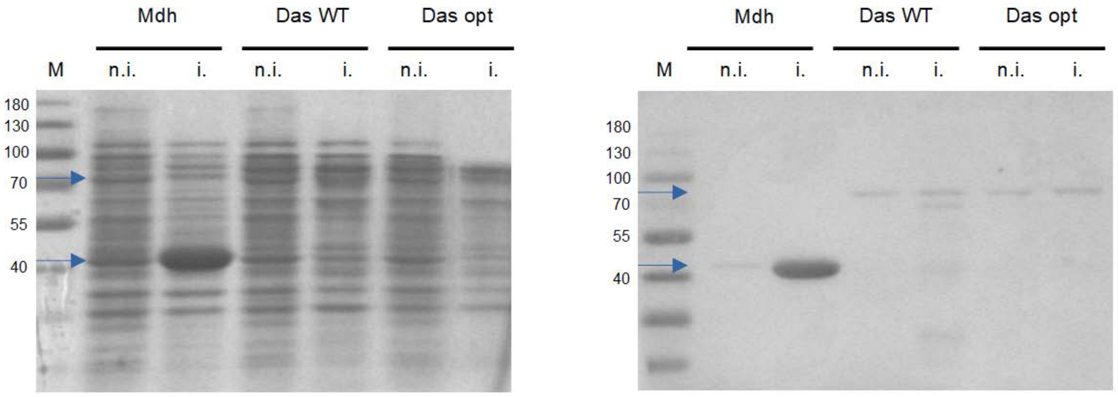


B)

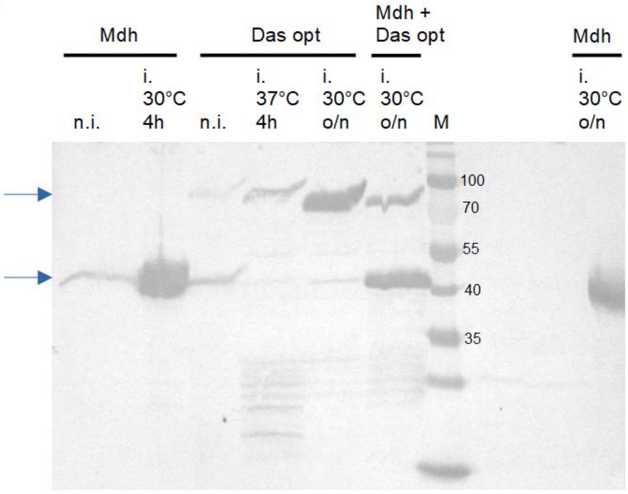


**Supplementary Figure S4. Expression analysis of *B. methanolicus* Mdh and *P. angusta* Das in different conditions.** (A) Cell lysates corresponding to $OD_{600}$ = 1.2 were separated on 12% SDS-PAGE gel and visualized by Coomassie staining (left) or Western blotted (right) using mouse anti-6×His antibody (1/10000) and anti-mouse AP-conjugated antibody (1/10000). Expression at 37°C for 4h after addition of 1 mM IPTG. M = molecular marker, n.i. = not induced, i. = induced. WT = wild-type, opt = codon-optimized. The expressed proteins are indicated by the blue arrows. Mdh = 40.8 kDa, Das = 78.7 kDa. (B) Western blot analysis of the expression at different incubation times and temperatures after IPTG induction (o/n = overnight).

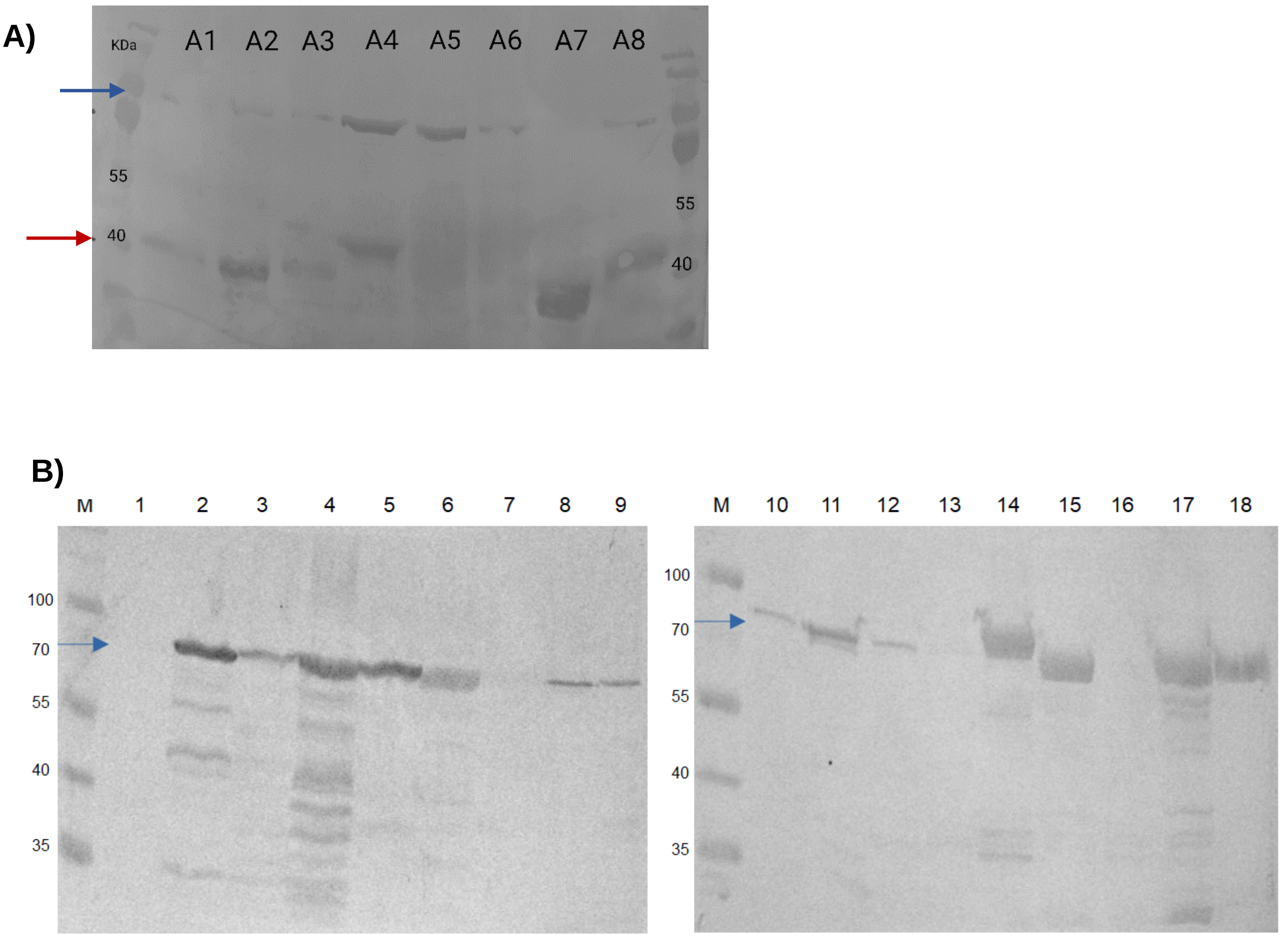


**Supplementary Figure S5. Western Blot analysis of expression of (A) Mdh and (B) Das homologues**. Cell lysates corresponding to OD = 1.25 were loaded. Induction with 1 mM IPTG was carried at 30°C overnight. The expressed Das are indicated by a blue arrow whereas the expressed Mdh are indicated by a red arrow. A) Mdh expression. A1: Bacillus methanolicus, A2: Pseudomonas lini, A3: Photobacterium damselae, A4: Bacillus coagulans, A5: Vibrio sp. ER1A (1), A6: Vibrio sp. ER1A (2), A7: Bacillus stearothermophilus, A8: Acinetobacter gerneri DSM 14967. The co-expressed Das is from P. angusta (opt). B) Das expression. 1: Aspergillus fumigatus, 2: Rasamsonia emersonii, 3: Exophalia acquamarina, 4: Diaporthe ampelina, 5: Aspergillus terreus, 6: Verruconis gallopava, 7: Kuraishia capsulata, 8: Fonseaca erecta, 9: Baudoinia compniacensis, 10: Magnoporthe oryzae, 11: Scedosporium apiospermium, 12: Pichia methanolica B (D7UPI3), 13: Debaryomyces hansenii, 14: Mycobacterium sp., 15: Pichia angusta (opt), 16: Dha reductase Pichia pfancensis, 17: Pichia angusta wild-type, 18: Pichia angusta wild-type not-induced. The codon-optimized version of Das (15) is more stable (no proteolytic degradation) compared to Das wild-type (17).

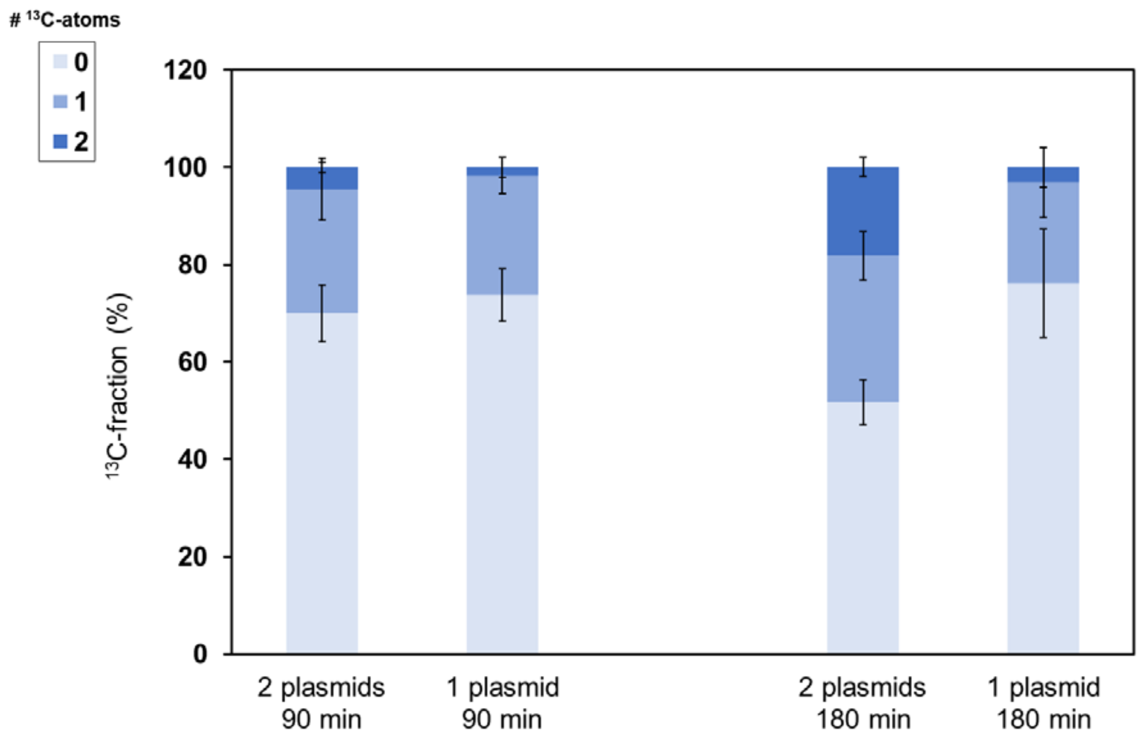


**Supplementary Figure S6. $^{13}$C-Methanol assimilation in the methylotrophic *E. coli* Δ*frmA* expressing the synthetic pathway from one or two vectors.** Labeling pattern of PEP after 90 min and 180 min of cultivation in M9 medium containing 655 mM $^{13}$C-methanol. Data presented as mean ± standard error of the mean (n = 3).

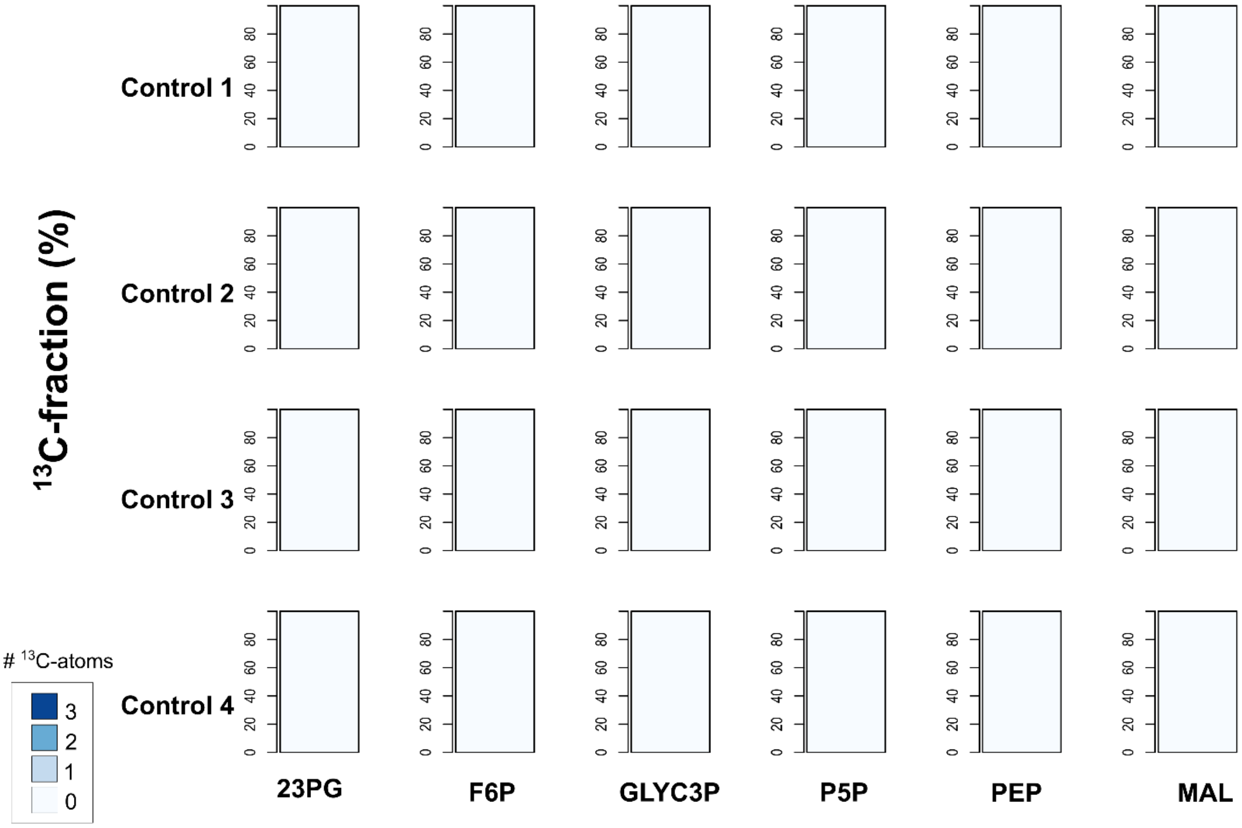


**Supplementary Figure S7: $^{13}$C-Methanol assimilation into central metabolism intermediates in the control strains of the genealogy of methylotrophic *E. coli*.** Labeling pattern of the intracellular metabolites 2 & 3 phosphoglycerate (23PG), fructose-6-phospohate (F6P), glycerol-3-phosphate (GLYC3P), pool of pentoses-5-phosphate (P5P), phosphoenolpyruvate (PEP) and malate (Mal) within the different control strains containing an empty plasmid after 90 min of cultivation in M9 medium containing 655 mM $^{13}$C-methanol.

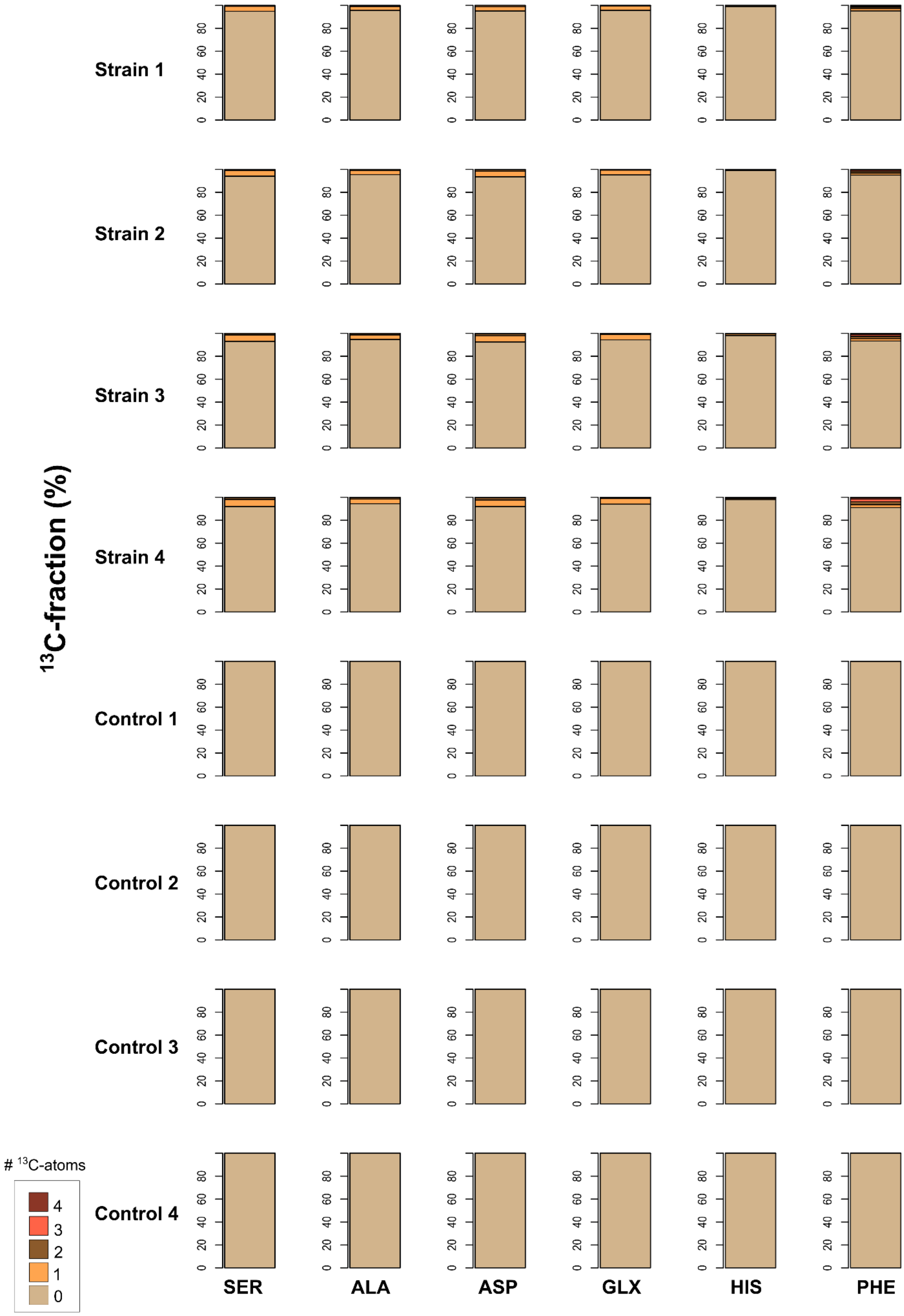


**Supplementary Figure S8: $^{13}$C-Methanol assimilation into proteinogenic amino acids in the genealogy of methylotrophic *E. coli*.** Labeling pattern of the proteinogenic amino acids serine (SER), alanine (ALA), asparagine (ASP), pool of glutamate and glutamine (GLX), histidine (HIS) and phenylalanine (PHE) within the different strains (see Figure 4B for details) and their respective controls containing an empty plasmid after 48h of cultivation in M9 medium containing 655 mM $^{13}$C-methanol.
